# Label-Free Quantification of Microtissue Growth Dynamics Using Optical Flow and Mitosis Detection

**DOI:** 10.64898/2026.08.10.743847

**Authors:** Kai Lennard Fastabend, Tassilo von Trotha, Konstantin Wolf, Rayan Chatt, Mario C. Benn, Viola Vogel, Philip Kollmannsberger

**Affiliations:** Biomedical Physics, Heinrich Heine University Düsseldorf, 40225 Düsseldorf, Germany; Laboratory of Applied Mechanobiology, Department of Health Sciences and Technology, ETH Zurich, Gloriastrasse 37/39, Zurich 8093, Switzerland; Department of Biotechnology and Biophysics, Biocenter, University of Würzburg, Am Hubland, 97074 Würzburg, Germany

**Keywords:** Microtissues, Optical Flow, Tissue Mechanics, Mechanobiology, Cell Divisions

## Abstract

While geometric constraints shape tissue development, quantifying the resulting growth dynamics remains a central challenge in tissue engineering. Conventional methods often struggle to capture multi-scale kinetics without complex labeling or difficult single-cell tracking. Here, we analyze geometrically controlled growth of microtissues derived from human dermal fibroblasts using time-resolved, label-free brightfield microscopy, combined with optical flow and semi-automated deep learning mitosis detection. By extracting multi-scale flow fields and integrating them with tissue segmentation, we quantify directional tissue dynamics, separating flow into parallel and normal components relative to the local tissue contour. Applying this framework, we contrast the quiescent tissue interior with the advancing growth front where localized dynamics and cell proliferation drive expansion. Our results demonstrate that, compared to the bulk, the growth front exhibits higher fluctuations parallel to the tissue contour, positive mean normal flow, and significantly increased mitotic activity. Furthermore, evaluating flow divergence around mitotic events reveals distinct spatial behaviors: with the onset of mitosis, a contraction and subsequent expansion occurs in the vicinity of the dividing cells. Beyond the immediate cellular neighborhood, the broader regional dynamics remain consistent before and after mitosis onset, with net tissue expansion in proximity to the growth front and contraction within the tissue interior. By extracting continuous kinetic data from easily accessible, label-free brightfield imaging, this approach serves as a non-invasive, complementary tool for evaluating in vitro tissue morphogenesis and growth dynamics. This analytical framework can be expanded to study locally resolved tissue morphogenesis and growth kinetics in other microsystems, ranging from embryos to organoids.

**Statement of Significance:** Understanding how localized cellular forces drive tissue growth is critical for mechanobiology. However, mapping these dynamics traditionally requires complex, invasive fluorescent labeling. We present an accessible, label-free computational framework combining optical flow and deep learning-based mitosis detection to quantify continuous tissue kinematics directly from standard brightfield microscopy. Applying this to 3D microtissues, we reveal a distinct spatial coupling between cell division, local mechanical fluctuations, and directed tissue expansion at the active growth front. This non-invasive approach bridges the gap between singlecell mechanics and macroscopic morphogenesis, offering a versatile tool to monitor complex in vitro model systems–like organoids and bioengineered tissues–without disrupting their native state.

## Introduction

Mechanobiology plays an important role in tissue growth, but is often overlooked compared to biochemical signals. Current understanding emphasizes the influence of tissue curvature and substrate geometry on organization, growth, and homeostasis of tissue (1). The continuous biomechanical deformations and structural rearrangements of living tissue arise from the reciprocal feedback-interactions between contractile cells and extracellular matrix (ECM) (2, 3): Contractile forces generated by the cytoskeleton allow cells to probe the mechanical properties of the ECM, while simultaneously reorganizing, remodeling, and tensioning their local matrix environment. In turn, ECM mechanics and architecture feed back to regulate cellular behavior, including proliferation, differentiation, secretion and cell fate decisions (4). Engineered 3D cell culture models, such as microtissues and organoids, provide powerful platforms for investigating the underlying mechanobiological processes (5, 6). It has been shown for a variety of cell types and model systems that concave substrate curvature promotes tissue growth by modulating tissue surface tension through cell-generated contractile forces (7–12). Experiments with de novo grown microtissues in engineered clefts show that tensile forces at the growth front trigger the differentiation of fibroblasts to contractile, proliferating, *α* SMA-positive myofibroblasts at the curved tissue-medium interface, while fibroblasts in the tissue core adopt a quiescent state (10). Building on these findings, a recent study investigated the fine-tuned reciprocal interactions between cells and ECM during transition from this tensed, myofibroblast- and fibronectin-rich growth front to a matured matrix with load-bearing collagen fibers populated by resting fibroblasts. Perturbing only one of the involved actors leads to a major delay of this transition, highlighting the tight organization of this shift (13).

While traditional end-point assays offer valuable insights into tissue composition, time-resolved analytical methods are required to capture continuous growth dynamics. To continuously quantify functional mechanical behaviors, including local contractility and structural deformation, non-destructive, label-free techniques like Particle Image Velocimetry (PIV) have been successfully applied to map deformation patterns around single fibroblasts in 3D collagen matrices (14). Additionally, at the cellular level, it has been shown that cell division transiently deforms the immediate extracellular neighborhood, with mitosis inducing localized cycles of contraction and expansion (15). However, while these label-free kinematic approaches succeed at the single-cell level, quantifying the continuous biomechanical deformations (16) during growth across an entire tissue would enable a better understanding of how localized cellular forces drive macroscopic morphogenesis. Applying these concepts to de novo grown 3D microtissues requires mapping a complex superposition of spatial dynamics, including cellular migration, local deformation, and continuous growth, relative to a curved tissue contour. Despite recent advances linking scaffold geometry to macroscopic tissue shape (10, 11, 13), the challenge is to precisely localize the spatio-temporal growth dynamics that drive continuous tissue morphogenesis. While short-term mechanical tension fluctuations are known to drive tissue fluidization and stress relaxation by enabling local cellular rearrangements (17), it is not known how these dynamics impact the macroscopic development of engineered 3D microtissues. Furthermore, although the mechanical footprint of cell division has been investigated around isolated cells in 3D collagen matrices (15, 18, 19), the combined analysis of these localized mitotic events and continuous macroscopic tissue development remains largely unexplored. Consequently, we lack a fundamental understanding of how single-cell mechanics scale up to the tissue level. It remains unclear how local contraction-expansion cycles driven by mitosis integrate into the macroscopic development of the active growth front versus the quiescent tissue interior.

To bridge the gap between single-cell local dynamics and macroscopic tissue morphogenesis, we developed a framework that extracts continuous kinematic readouts from labelfree brightfield time-series data using a novel combination of multi-scale optical flow analysis and deep learning-based mitosis detection. We applied this approach to de novo grown microtissues in engineered cleft geometries (13). Human fibroblast cells, after covering the available free PDMS surface, start to form a dense three-dimensional fibrous tissue in the concave cleft regions. This model system was chosen because it provides a highly reproducible, spatially defined transition from an active, tension-driven growth front to a quiescent tissue interior, making it the ideal platform to map localized biomechanical behaviors. First, we employed the Farnebäck algorithm for optical flow (20) to calculate dense motion fields between consecutive frames by extracting pixel-by-pixel displacement vectors from local intensity changes. To relate these kinematics to the tissue geometry, optical flow vectors were locally separated into normal and parallel components relative to the contour of the microtissue growth front. Second, to link these tissue-scale mechanics to localized biological events, we trained a deep learning object detection model based on the YOLO framework (21) to assist the manual annotation of mitosis events. We selected this object detection architecture for its exceptional speed and robustness in identifying cellular events within large microscopy datasets.

Analyzing these combined readouts revealed distinct spatial patterns across the developing tissue: the active growth front was predominantly characterized by dynamic transients, defined as optical flow magnitudes that exceed baseline fluctuations. Correspondingly, local divergence of the flow fields showed consistent tissue expansion in this active growth front. By further separating the directional flow components relative to the tissue contour, we revealed increased parallel fluctuations and mean positive normal deformations within this expanding region. Furthermore, the temporal and spatial distribution of mitosis events directly complemented this kinematic data, showing a strong accumulation near the growth front that spatially correlated with the increased optical flow dynamics. Finally, a localized spatio-temporal analysis of flow field divergence directly around these detected events revealed distinct, transient contraction and expansion patterns associated with the mitotic activity.

Our methodological framework allows researchers to extract comprehensive kinematic data, such as local divergence and directional flow components, from standard brightfield microscopy data that is often routinely acquired for non-invasive monitoring, without relying on additional labeling. This approach complements existing techniques by correlating specific biological events, such as mitosis, with biomechanical parameters such as localized contraction and expansion. Our findings demonstrate that macroscopic tissue growth is orchestrated by spatially distinct zones of active deformation and cellular proliferation. The ability to continuously quantify fluctuations and expansion patterns at the active growth front compared to the quiescent tissue interior provides a label-free readout for evaluating localized tissue formation. By linking the spatial occurrence of cell divisions with localized kinematic changes, this work bridges the gap between single-cell behavior and global tissue mechanics.

## Materials and Methods

### Fabrication and functionalization of substrates

Fabrication and functionalization of Polydimethylsiloxane (PDMS) substrates followed previously established protocols (13). Briefly, microstructures were patterned on 4-inch silicon wafers using photolithography with multiple layers of negative photoresist (SU8 3050, micro resist technology). PDMS was prepared using the Sylgard 184 Silicone Elastomer Kit (Dow Chemical Company) by mixing at a 10:1 base-to-curing agent ratio. Scaffolds were cast from the SU8 master using a custom-built squeezer device that removed excess liquid PDMS during curing, producing thin PDMS sheets containing V-shaped clefts (45°) as defined by the SU8 microstructures. The PDMS in the compression device was cured for at least 18 h at 60°C before being punched into 18 mm circular substrates. These substrates were treated with 1 mg/ml sulfo-SANPAH (22589, Thermo Fisher Scientific) in PBS (pH 8.5) and exposed to UV light for 150 s (365 nm, 16 mW/cm^2^; RX FireFly 75×20AC365, Phoseon Technology), followed by incubation with 20 µg/ml fibronectin in PBS (pH 7.4) for 1 h at room temperature to enable covalent binding. The fibronectin used in this step was isolated from human plasma (Blutspende SRK Zürich) according to an established protocol by Speziale et al (22). The functionalized substrates were then mounted into dishes with a polymer bottom (µ-Dish 35 mm; 81151, ibidi) using UV-cured NOA-61 glue (C007030-6, Norland). To prevent unwanted cell adhesion, the dish bottom was passivated with mg/ml poly-L-lysine (20 kDa) grafted with poly-ethylene glycol brushes (5 kDa; PLL(20)-g[3.5]-PEG(5), SuSoS). Assembled scaffolds were sterilized by UV for 30 min, washed with PBS, and equilibrated in culture medium at 37 °C for at least 30 min prior to cell seeding. A schematic overview of the V-shaped substrate geometry and the resulting 3D microtissue morphology following cell seeding is illustrated in Fig. 1 A.

**Fig. 1.**
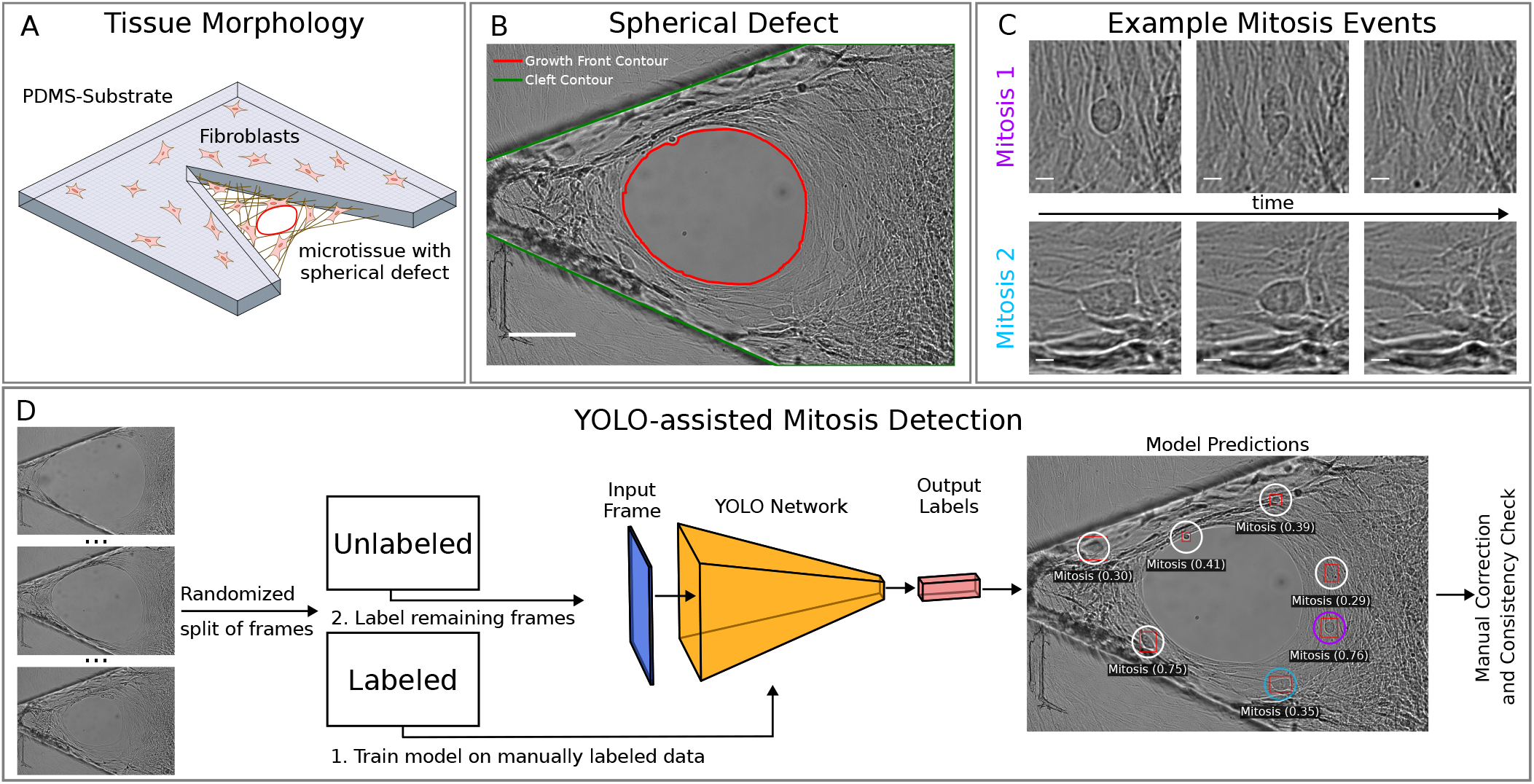
YOLO-assisted detection of mitosis events in microtissue time series. **(A)** Schematic illustrating the V-shaped PDMS substrate and the resulting 3D morphology of the microtissue grown within it. The flat surface of the PDMS is covered with a confluent fibroblast monolayer, while the 3D microtissue is anchored to the cleft walls. The tissue shows a circular defect resulting from a central rupture, which closed during subsequent growth. **(B)** Representative bright field image of the microtissue time series with segmented cleft and growth front contour (scale bar: 100 µm). The growth front (red) is defined by the shape of the circular defect. **(C)** Temporal development of two exemplary mitosis events over three consecutive frames (frame interval: 639 s; scale bar: 10 µm). **(D)** Schematic of the YOLO-assisted mitosis detection pipeline. The frames of the time series are divided into two parts at random. One part of the frames are labeled manually and used to train a YOLO network for mitosis detection. Subsequently, unlabeled frames are passed through the trained model to predict mitosis events. The predictions are filtered by confidence scores (confidence threshold: 0.2) and displayed with red bounding boxes. Finally, the predicted labels are corrected manually and checked for consistency with the labeled frames used for training. Overlapping bounding boxes in consecutive frames are assigned to the same mitosis event. Purple and blue events in the model predictions correspond to the events shown in (C).

### Cell culture

Primary normal human dermal fibroblasts (NHDFs; C-12300, PromoCell) were cultured in αMEM (Biowest, L0475) supplemented with 10% fetal bovine serum (VWR, S181H-500) and 1% penicillin-streptomycin (Thermo Fisher Scientific, 15140122). For microtissue experiments, cells were detached using 0.05% Trypsin-EDTA (25300054, Thermo Fisher Scientific) and seeded at a density of 2 × 10^5^ cells per functionalized substrate. From the time of seeding, the culture medium was supplemented with 100 µM L-ascorbic acid (A8960, Sigma-Aldrich) at each medium change. Cultures were maintained at 37°C in a humidified atmosphere containing 5% CO, and the medium was replaced every 2-3 days.

### Microscopy

Brightfield time-lapse imaging of microtissue growth was performed using a Zeiss Axio Observer 7 equipped with a 20x air objective (0.4 NA, LD Plan-Neofluar Korr Ph2). During image acquisition, cells were maintained in a humidified atmosphere at 37°C with 5% CO2. A representative image from the resulting time-lapse series is shown in Fig. 1 B.

### Farnebäck Algorithm for Optical Flow Motion Estimation

To calculate the displacement map between two consecutive frames in a time series, we employed the Farnebäck algorithm for dense optical flow (20). The core concept of this algorithm is to treat the pixel intensities of an image as a 2D surface or landscape. Locally, this surface is approximated by a second-order polynomial, which captures the local intensity gradients and curvatures in its linear and quadratic terms. The polynomial coefficients are estimated over a spatial neighborhood via weighted convolutions to increase noise robustness, with a trade-off in spatial resolution. Details on the mathematical background are given in Supplementary Note 3. Assuming that local brightness remains constant between consecutive frames, the algorithm calculates motion by determining how far, and in what direction, these local polynomial surfaces have shifted. This method calculates a motion vector for every single pixel, creating a dense motion field.

Because the algorithm tracks entire local patches of pixels rather than isolated points, the resulting motion field is inherently smooth, naturally preventing abrupt, noisy shifts between neighboring pixels. Reliable motion estimation requires sufficient local image texture, defined by spatial intensity variations, such as distinct edges, corners, or patterns. In locally flat or edge-dominated regions, the polynomial representation might not contain enough independent information to reliably determine motion in all directions, a fundamental limitation known as the aperture problem.

The estimation of motion relies on small inter-frame displacements relative to the size of the spatial neighborhood used to approximate the local image intensity. The detection of larger motion is typically handled by iterative refinement on different spatial scales. The implementation in OpenCV (23) performs this type of pyramidal processing with increasingly finer resolution, whereby the motion estimation of the previous, coarser scale is refined iteratively. Next to the parameters for number of pyramids and scale, there are a number of parameters that effect the amount of spatial smoothing of the resulting flow field. At the core of the parameter choice, a balance must be struck between the possible spatial resolution and the sensitivity for fast large scale motion. A detailed list of the used parameters is given in Table S4. Using eq. 1, the local divergence (Fig. 2 D) of the vector field can be calculated, to analyze the spatial distribution of sources and sinks of the optical flow field. Assuming that the flow field is dominated by deformations in the tissue, positive divergence represents regions of tissue expansion, whereas negative divergence corresponds to tissue compression.

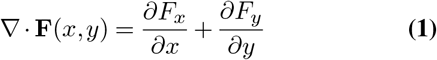

**Fig. 2.**
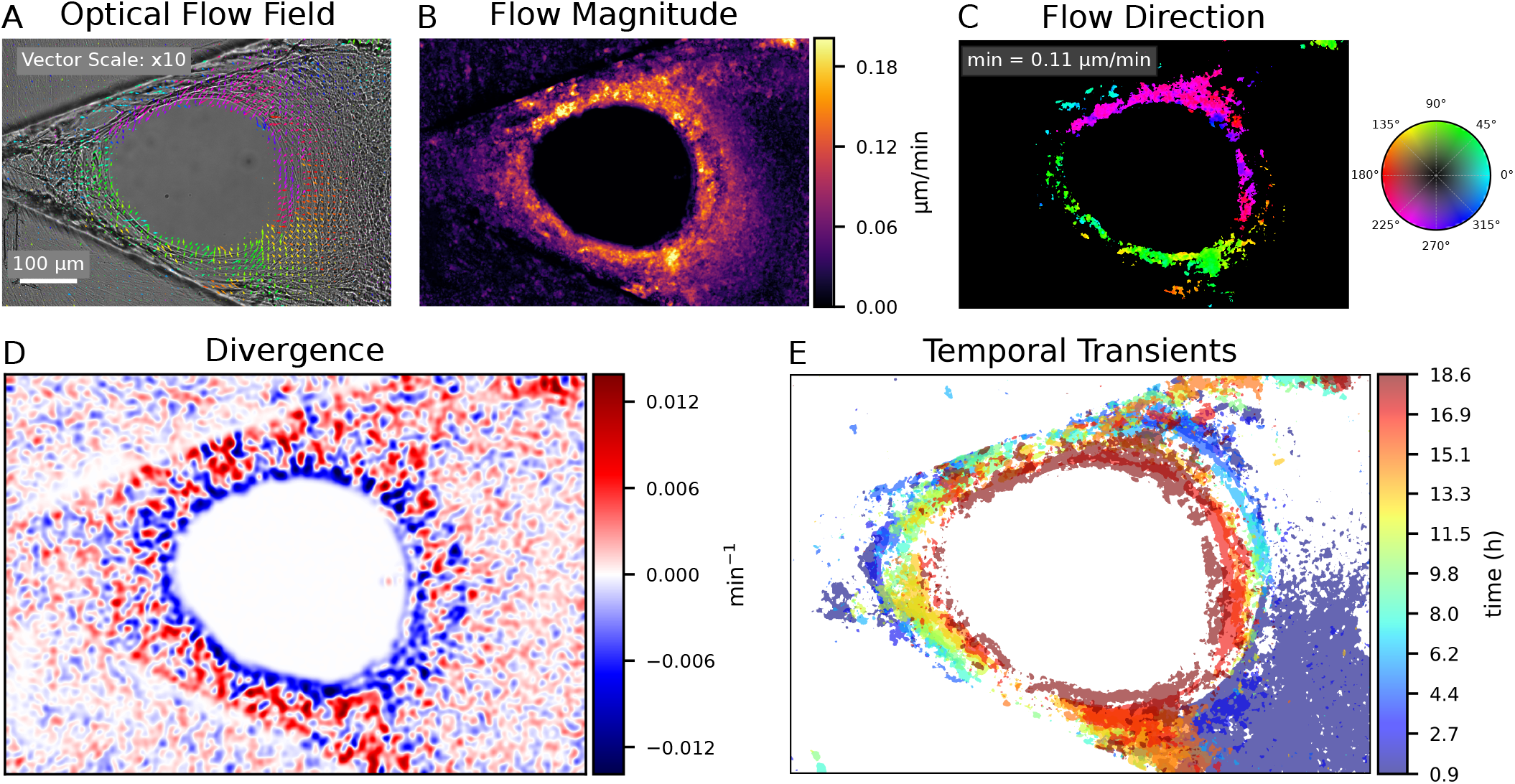
Optical flow analysis reveals localized dynamics during the closure of a microtissue defect. The analyzed microtissue, originally grown in a V-shaped cleft, experienced an internal rupture that led to a circular defect. The analysis highlights the tissue dynamics during the subsequent closure process of this defect. **(A)** Representative optical flow field, visualized as a vector field on the first image of the considered time window (stride: 50 px; vector scale: x10). Flow fields are averaged over 10 consecutive frames to highlight persistent long-term motion. **(B-C)** The flow magnitude (B), which represents the local speed of the optical flow, and direction (C) illustrate the pixel-level information contained in the flow field. The flow direction is filtered by a minimum velocity threshold of 0.11 µm/min to isolate significant dynamic transients. **(D)** Divergence of the optical flow field: positive divergence (sources; red) indicates regions of apparent tissue expansion, while negative divergence (sinks; blue) corresponds to apparent compression. The divergence map reveals granular spatial patterns of alternating expansion and compression, highlighting pronounced flow convergence at the growth front. **(E)** Temporal evolution of dynamic transients (> 0.11 µm/min) calculated across non-overlapping time windows for the full duration of the time series, color-coded by time. The spatial distribution of the temporal transients coincides with the progressively closing pore, illustrating the spatio-temporal dynamics of the growth front.

The divergence of the flow field was computed as the sum of second-order central differences. To extend the analysis to the cellular scale and capture large-scale divergence patterns, the input fields were spatially smoothed using a Gaussian blur with a kernel size of 101 px (*σ* = 15.5), effectively integrating neighborhood information and ensuring the divergence values reflect biological motion on the scale of individual cells. While this facilitates robust tracking at the cellular scale, it inherently involves a trade-off in spatial resolution; highly localized, sub-cellular divergence patterns are attenuated, and discrete motion events may be spatially broadened into surrounding areas.

### YOLO Object Detection

Deep learning-based computer vision algorithms are increasingly applied to biological datasets to automate complex visual tasks, such as cell tracking and the recognition of dynamic cellular events. To locate and classify characteristic mitosis events (Fig. 1 C) within our image time series, we employed the YOLOv5 object detection framework (24). YOLO formulates object detection as a single-stage regression problem, allowing for highly efficient bounding box and class probability predictions (21).

The standard network architecture and tensor outputs are extensively documented in the literature (24, 25). Therefore, we focus solely on the specific application of the model to mitosis detection. The model predicts candidate bounding boxes characterized by spatial coordinates, dimensions, as well as confidence and objectness scores. These outputs allow the network to effectively discriminate between regions containing mitotic cells and empty background.

To minimize the time-intensive effort of manual annotation, we implemented a YOLO-assisted semi-automated workflow (Fig. 1 D). Initially, a randomized 50% subset of the time series frames was manually annotated using *LabelImg* (26), identifying cells undergoing mitosis based on their characteristic rounded morphology. This dataset was split into 80% for training and 20% for validation to ensure robust generalization across different time points.

We utilized the pretrained YOLOv5s model variant provided by Ultralytics (24). The model was fine-tuned for 100 epochs using a Stochastic Gradient Descent (SGD) optimizer. Training was performed on images resized to 1280 pixels with a batch size of 2 and high-augmentation hyperparameters (detailed in Table S2). Model performance was evaluated using standard computer vision metrics, including precision, recall, F1-score, and mean Average Precision (mAP). To ensure high-precision spatial localization, the “best weights” for the final model were selected from the training epoch that achieved the highest mAP@0.5:0.95. This rigorous metric averages the mAP across a range of Intersection over Union (IoU) thresholds, effectively penalizing poor bounding box alignment.

During inference, the trained model automatically generated bounding box predictions for the remaining unlabeled frames. These predictions were filtered using a deliberate, low confidence threshold of 0.2. While this lower threshold increases initial false positives, it crucially minimizes missed mitosis events; the subsequent manual correction phase remains significantly faster than finding and annotating overlooked events from scratch. Finally, overlapping bounding boxes in consecutive frames were linked to the same mitosis event, and all YOLO-assisted labels underwent a final manual consistency check against the initial annotations.

### Tissue Segmentation

The workflow for the automated segmentation of the tissue contour is shown in Fig. S2. Initially, a Sobel-Filter for edge detection was applied and the resulting gradient image was smoothed with a Gaussian to highlight large scale edges. Afterwards a binary mask was defined using a threshold and the mask was inverted and smoothed. A final erosion step of the mask yielded a contour that corrects for an offset from the actual tissue contour in the image due to previous smoothing steps. The used parameters for the segmentation steps are given in Table S3.

### Curvature Calculation

To quantify the morphological changes of the tissue boundary, we required a method robust against the pixel-level jaggedness of digital segmentations. Drawing conceptually on the scale-dependent approach of Bidan et al. (27, 28)–though utilizing a distinct geometric implementation–we evaluated the boundary over a macroscopic neighborhood rather than using immediately adjacent pixels. Specifically, the signed curvature *κ* was estimated for each contour point *P*_0_ via a three-point circumcircle approximation (29). By selecting two neighboring points *P*_−_ and *P*_+_ at a fixed contour distance of *s* = 50 px, we defined a unique triangle that approximates the local osculating circle while intrinsically filtering high-frequency noise.

The magnitude of the curvature is given by the inverse of the circumradius of this triangle:

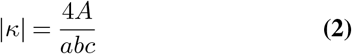

where *a, b*, and *c* represent the Euclidean distances between the three points *P*_−_, *P*_0_, and *P*_+_. The triangle area *A* was computed using the 2D cross product of the vectors connecting these points:

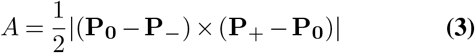

The sign of the curvature was determined by the orientation of the turn, derived directly from the sign of the cross product, where *κ* > 0 indicates a convex (left) turn and *κ* < 0 denotes a concave (right) turn.

### Computational Hardware and Processing Time

All computational analyzes, including optical flow estimation and YOLO model training, were performed locally on a laptop workstation running Windows 11 Pro. The system was equipped with a 13th Gen Intel Core i7-13850HX processor, 64 GB of RAM, and an NVIDIA RTX A1000 Laptop GPU with 6 GB of VRAM. Under these hardware specifications, the computation of the dense optical flow fields using the Farnebäck algorithm required approximately 1.4 seconds per frame, totaling less than 3 minutes for a representative 122-frame time series. Fine-tuning the YOLOv5s model for 100 epochs on the annotated dataset took approximately 7 minutes.

## Results

### Optical Flow Fields Capture Increased Growth Front Dynamics

To quantify continuous multi-scale tissue kinetics, we applied an optical flow analysis to our time-resolved microscopy data to determine the spatial distribution of tissue dynamics. The microtissues analyzed in this study were grown in V-shaped clefts, where an internal rupture led to a circular defect that progressively closed during growth (see Fig. 1 A and B). The main focus of this analysis is to evaluate the tissue dynamics during the closure of this defect, which specifically enabled us to distinguish the active growth front from the stabilizing mature tissue. Figure 2 A shows an exemplary optical flow field averaged over 10 frames of the tissue time series, which serves as a mean field highlighting consistent long-term dynamics in the respective time window. The flow field magnitude shown in Fig. 2 B displays the local speed of the captured dynamics at pixel resolution. Figure 2 C shows the corresponding flow directions of filtered dynamic transients–defined as local optical flow magnitudes that overcome baseline fluctuations, thereby highlighting regions with the fastest motion–that overcome a speed of 0.11 µm/min. The exemplary mean field reveals a granular structure of the motion field and enables the identification of coherent movement patterns at high spatial resolution. The field shows localized transients in proximity to the growth front, which decrease in deeper layers of the maturing tissue. Furthermore, there is a clear difference in flow field magnitude between the cleft interior and the flat substrate.

The divergence of the optical flow vector field highlights local sources and sinks, corresponding to diverging or converging motion patterns. As shown in Fig. 2 D, it reveals a granular structure of regions with apparent contraction (blue) and expansion (red). Interestingly, the tissue contour of the pore is characterized by a circular region with negative divergence, indicating a net converging motion close to the growth front. In deeper tissue layers, however, expanding and contracting regions are randomly interspersed, displaying a heterogeneous distribution without a distinct macroscopic pattern. Biologically, this localized contraction aligns with the known presence of *α*-SMA-positive cells at the growth front (13), demonstrating that our brightfield analysis can identify the contractile growth front without requiring immunofluorescence staining.

To visualize the development of dynamic transients over the course of the time series, transients of consecutive time windows are overlayed in Fig. 2 E. In addition to the description of the spatial distribution in a single flow field, the localization of the dynamic transients near the closing defect contour is once again evident here. The temporal color gradient of the transients indicates the layered spatial distribution of the successive layers of tissue, matching experimentally observed layered growth patterns (30). This highlights the connection between the closing pore contour and the transients near the growth front, which are spatially correlated. The threshold for the temporal transients was determined based on the results of the dynamic fluctuations determined in Fig. 3 to filter out optical flow dynamics with a low magnitude.

**Fig. 3.**
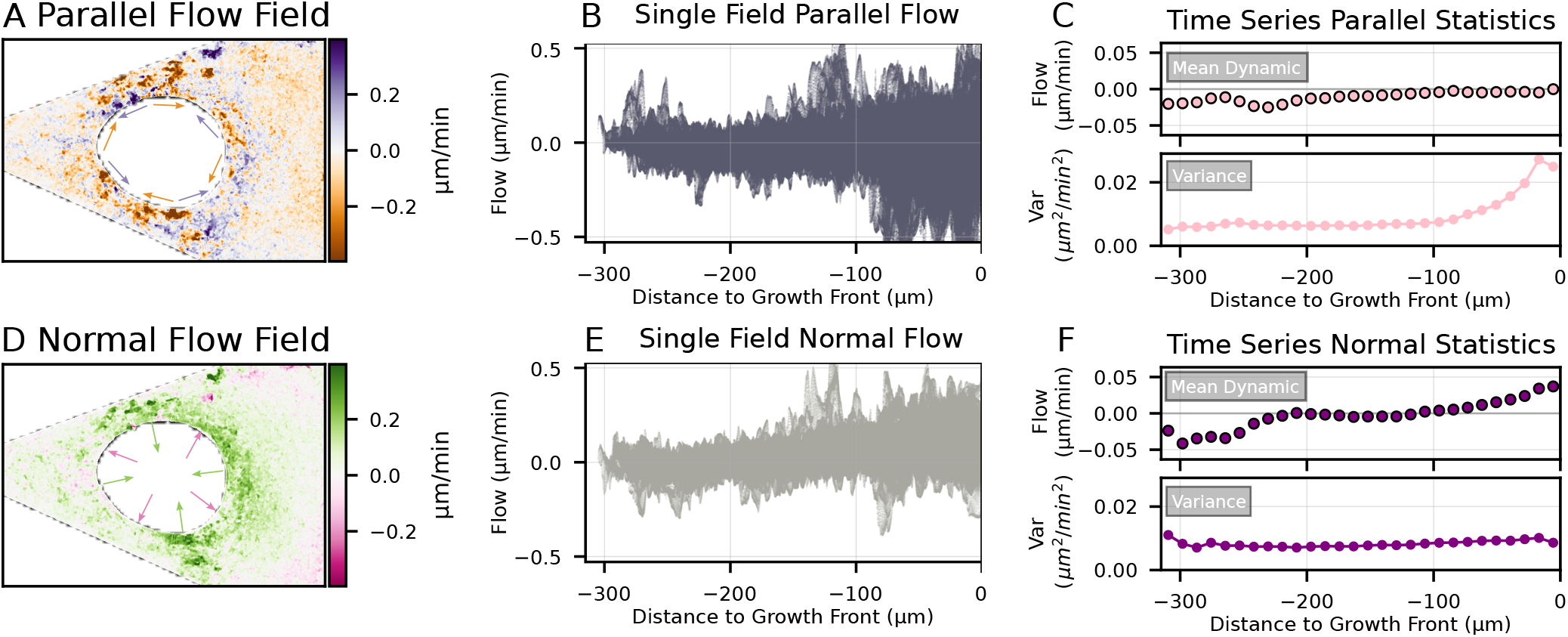
Geometric quantification of normal and parallel flow relative to the growth front located at the edge of the circular defect. **(A,D)** Local optical flow separated into parallel (A) and normal (D) components relative to the growth front contour, visualized as scalar fields. Arrows indicate the local direction for each component at the tissue contour, corresponding to the diverging color scales. **(B,E)** Scatter plots of local magnitude against growth front distance for parallel (B) and normal (E) flow of a single field. Each point in the scatter plot corresponds to one pixel in the scalar field. **(C,F)** Temporal statistics of mean flow and variance over the entire time series, calculated in 50 px distance bins. Parallel statistics (C) reveal increased fluctuations and zero-mean flow in proximity to the growth front, while normal statistics (F) are characterized by positive mean dynamics at the growth front with consistently low variance across the entire tissue.

In addition to the static snapshots in Fig. 2 B and C, Supplementary Movies 1 and 2 provide the continuous evolution of flow direction and magnitude over time at pixel resolution. Furthermore, Fig. S3 shows supplementary data, analogous to the displayed pore tissue (Fig. 2), for early-stage growth of tissue in the tip of the cleft. While the additional data confirms the key observations regarding dynamic transients near the growth front, the associated negative divergence is noticeably less pronounced than at the circular defect. Biologically, this can be attributed to the distinct shape of the early-stage tissue. A shorter growth front with reduced mean curvature likely generates less positive mechanical feedback among the contractile cells compared to the closed, circular defect.

### Flow Components Reveal Parallel Fluctuations and Normal Tissue Expansion at the Growth Front

Figure 3 shows the analysis of separated parallel and normal components relative to the contour of the circular defect. The underlying component separation is illustrated in detail in supplementary Fig. S4. In brief, this method calculates the shortest distance from each pixel to the contour to define a local normal vector, which is then used to decompose the local tissue flow into its normal and parallel components. The resulting components are displayed as color-coded spatial heatmaps in Fig. 3 A for the parallel flow and in Fig. 3 D for the normal flow. The local directions of positive and negative flow are indicated by arrows matching the respective color maps. The parallel flow field reveals prominent transients in proximity to the growth front–defined here as the advancing inner edge of the circular defect–with opposing regions of parallel movement along the defect’s contour. Compared to the parallel component, the normal flow field shows mainly positive dynamics oriented toward the growth front, with negligible regions exhibiting negative normal deformation for the considered time window. This indicates tissue movement that actively contributes to the closure of the circular defect.

Parallel and normal dynamics vary distinctly with distance from the growth front (Fig. 3 B, E); while the parallel distribution scatters widely around zero, normal dynamics tightly cluster with a positive mean shift near the front. To systematically evaluate the long-term flow characteristics, Fig. 3 C and F present mean and variance statistics from cumulative scatter plots of parallel and normal dynamics covering the entire time series. Since different distance intervals from the growth front contain varying numbers of pixels, we analyzed the variance and mean values in curved distance bins (11.25 µm wide) that follow the contour of the growth front, thereby normalizing for differences in contributing tissue area.

Analyses of the flow dynamics reveal that the variance of the parallel component is non-uniformly distributed relative to the distance to the growth front, manifesting as two distinct spatial zones: a highly dynamic region within the first 100 µm characterized by strong positive and negative peaks, followed by a transition into a lower-variance regime deeper in the tissue. In comparison, the variance of the normal dynamics maintains a consistent level independent of the distance to the growth front. The mean-zero values of the parallel dynamics near the defect edge show that the parallel flow has no preferred direction and that these strong fluctuations are symmetrical over the course of the time series. However, the normal component exhibits positive mean values that also fall off after 100 µm. This reveals consistent long-term dynamics driving the closure of the defect pore. Interestingly, deeper layers of tissue (around 250 µm) show negative normal flow. This could be related to the out-of-frame growth front of the tissue on the right side of the image, which is consistent with the negative signal in Fig. 3 D. The contour of this additional growth front, advancing towards the cleft opening, is comparable to the early-stage tissue shown in Fig. S3 and has been extensively characterized in previous work by Benn et al. (13). By geometrically decoupling the flow components, this framework effectively separates symmetrical, mean-zero fluctuations from directed expansion kinetics, providing a powerful, label-free method to map the precise mechanical transitions that drive tissue morphogenesis.

### YOLO Accelerates Manual Annotation of Mitotic Events

In label-free brightfield microscopy, manually tracking mitotic events within dense microtissues is a significant bottleneck. To overcome this, we integrated a YOLO-assisted mitosis detection workflow (Fig. 1 D) as an assistive tool. Because our primary objective is the reliable identification of mitotic events to extract temporal dynamics rather than strict spatial bounding box localization, we fine-tuned a YOLO model specifically to handle the visual features of our datasets. To evaluate its performance and estimate potential time savings, we quantified the manual labeling time versus the time required to correct the YOLO labels using a representative data set. While baseline manual annotation required approximately 60 seconds per frame, correcting the proposed labels of the fine-tuned YOLO model required only a fraction of that time (see Table S1). When the actual training duration, automated inference, and final consistency checks are factored in, the YOLO-assisted workflow leads to an overall 53% decrease in annotator time compared to fully manual annotation. Crucially, this provides the additional advantage that training and inference are computationally automated, freeing up active researcher time.

The results of the YOLO training on the manually annotated frames are shown in Fig. S1. The rising mean average precision values (mAP@0.5 and mAP@0.5:0.95) indicate the learning progress of the YOLO model over the 100 training epochs. The object and box losses for the training and validation data decrease in parallel, indicating no strong overfitting and confirming that the model generalizes well to the validation data. The best mAP@0.5:0.95 = 0.28 is reached in epoch 85 and was used to define the best weights of the trained model. Notably, the model achieves a robust mAP@0.5 = 0.68 at this epoch, highlighting a high overall detection quality that successfully satisfies the requirements for our semiautomated annotation pipeline.

### Mitotic Activity is Increased in Proximity to the Curved Tissue Contour

Having established that the growth front exhibits distinct mechanical signatures–specifically, strong parallel fluctuations and net normal expansion–we hypothesized that these localized physical dynamics are driven by underlying biological activity. To investigate the cellular origin of this tissue expansion, we correlated the analysis of mitosis events derived from our YOLO-assisted labeling with the analysis of macroscopic tissue properties, like area and curvature, extracted via automated tissue segmentation Figure Qualitatively, the spatial distribution of these proliferation events appears to track the closing growth front, with later mitosis events (highlighted in red) clustering closer to the inner pore contour (Fig. 4 A). This layered progression of the advancing tissue boundary over time is explicitly mapped in Fig. 4 B, illustrating the gradual closure of the defect. By evaluating active mitosis events relative to the tissue contour of the exact corresponding time point, we extracted the spatial distribution of mitosis events as a function of distance from the growth front (Fig. 4 C). The absolute count of mitosis events reveals highly localized mitotic activity in close proximity to the advancing growth front. Notably, this spatial profile has a strong correlation with the distribution of parallel fluctuations and mean-positive dynamics measured with optical flow (Fig. 3 C and F). While the physical expansion and fluctuations fall off after a growth front distance of 100 µm, the main peak of the mitosis distribution drops on a nearly identical spatial scale, dropping significantly after 75 µm. This demonstrates a clear spatial overlap between the physical expansion and fluctuations captured by the flow fields and the regions of elevated cellular proliferation.

**Fig. 4.**
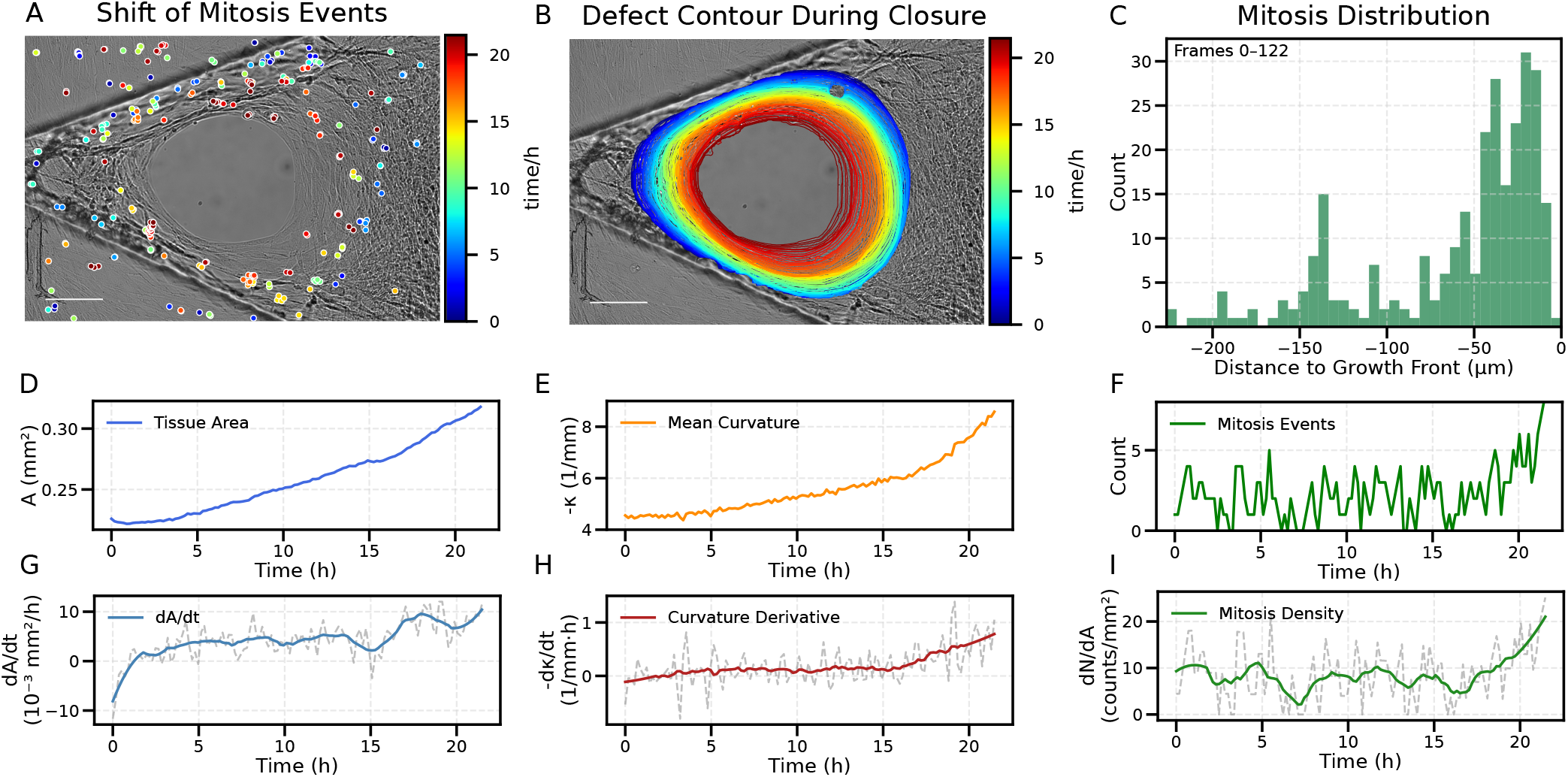
Analysis of mitosis events and tissue morphology. **(A)** Spatio-temporal distribution of detected mitosis events superimposed on the final frame of the time series. **(B)** Temporal evolution of the defect contour, illustrating the gradual closure and progression of the growth front. Contours and mitosis events colored the same in A and B correspond to the same point in time in the time series (shared scale bar: 100,µm). **(C)** Spatial distribution of mitosis events relative to the growth front distance during defect closure. Mitosis events were binned according to their distance from the instantaneous tissue contour (bin size: 40 px). The resulting distribution reveals a localized accumulation of mitosis events in close proximity to the growth front, with a significant decay beyond 75 µm into the mature tissue. **(D–I)** Kinetics of tissue growth and mitotic activity. **(D, G)** Evolution of microtissue area within the cleft and its corresponding growth rate. **(E, H)** Mean contour curvature and its rate of change over time. **(F, I)** Total count of active mitosis events and area-normalized mitosis density. Metrics in (G, H, I) represent smoothed temporal derivatives to highlight long-term trends.

To determine whether this spatial correlation translates into a temporal coupling between tissue morphogenesis and cellular proliferation, we tracked the kinetics of growth, curvature, and mitotic activity over the course of the time series (Fig. 4 D–I). The analyzed tissue area is defined as the tissue inside the cleft, excluding the area of the open pore. A comparison between the tissue area and the mean curvature of the tissue contour reveals similar trends, characterized by a nonlinear increase in both metrics that accelerates toward the end of the measurement period. The time derivatives of both metrics highlight that tissue growth and curvature increase in parallel. Interestingly, the growth rate deviates from a strict monotonic increase and exhibits temporal fluctuations that correlate with the temporal course of the normalized mitosis density. Overall, the macroscopic geometric metrics exhibit tightly coupled temporal profiles; however, their correlation with mitotic density remains weak, becoming detectable only in the final stages of the time series.

### Divergence Around Mitosis Events Reveals Regions of Tissue Contraction and Expansion

To further investigate the observed correlation between mitotic activity (Fig. 4 C) and tissue dynamics in proximity to the growth front (Fig. 3 C and F), an analysis of the mean divergence was performed around the detected mitosis events using segmented radial regions of interest (ROIs). We adapted these radial sectors from previous work mapping localized fibroblast contractions in collagen (31). Because our prior analysis demonstrated that the main peak of mitotic activity sharply declines beyond 75 µm from the defect edge (Fig. 4 C), we restricted this localized analysis entirely to events occurring within the active growth front (N = 90). Figure 5 A shows a representative mitosis event overlaid with ring-shaped ROIs separated into segments with an angle interval of 45°. Relative to the normal vector pointing toward the growth front, the 0°-Segments are directed toward the defect edge, while other segments are ordered clock-wise. The radii 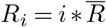 were defined by the mean bounding box size 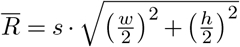 of all considered mitosis events, such that the central region represents the dividing cell, while the surrounding rings correspond to progressively larger interaction ranges. Overlapping detections in consecutive frames were assigned to a single mitotic event to enable dynamic analysis in the temporal context before and after mitosis onset. The first frame that belongs to one event is referred to as the start of mitosis.

**Fig. 5.**
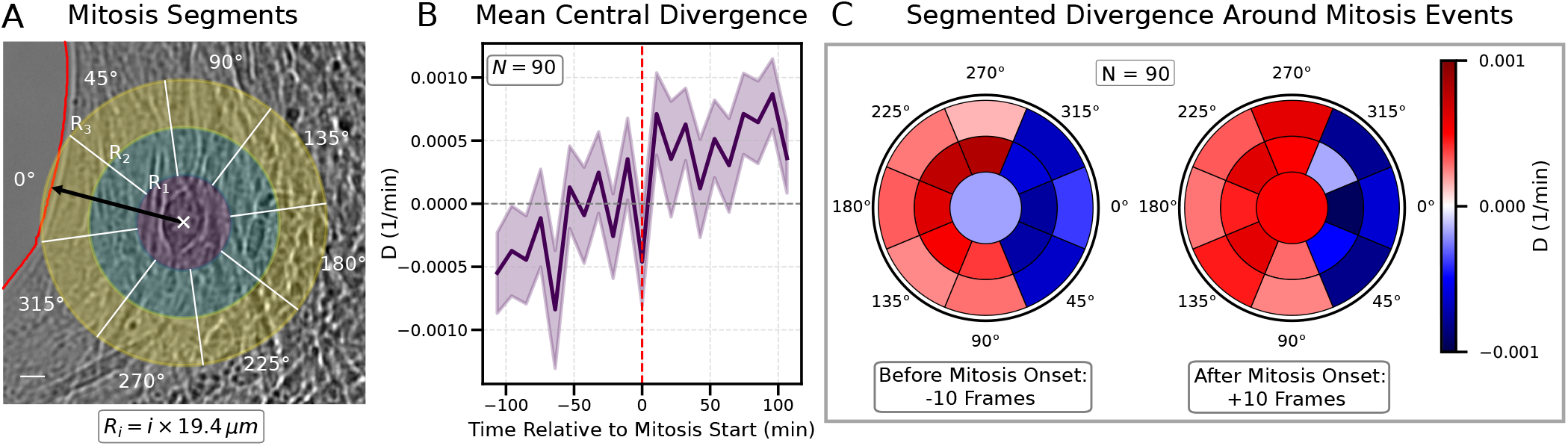
Spatio-temporal divergence patterns surrounding mitosis events. In this analysis, positive divergence corresponds to apparent local tissue expansion, while negative divergence indicates local tissue compression (convergence). **(A)** Representative mitosis event with overlaid segmented regions of interest (ROIs). A central circular ROI is surrounded by concentric rings, partitioned into 45° angular sectors. The 0° sectors are oriented toward the growth front, while 180° sectors point toward the tissue interior. Ring radii are defined as 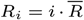, where 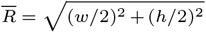 is the mean size of all detected mitosis bounding boxes. Only events within 75 µm of the growth front were included in the analysis (*N* = 90) **(B)** Mean divergence of *N* = 90 events within the central ROI over a window of *±* 10 frames relative to mitosis start. The data exhibit a sign reversal, transitioning from convergent dynamics before mitosis to divergence after the event start. **(C)** Polar plots of mean divergence (*N* = 90) relative to the tissue contour, averaged over 10 frames before and after mitosis start. While the central region undergoes a sign reversal, the surrounding rings reveal a consistent macroscopic pattern: tissue convergence toward the growth front (0°) and expansion toward the interior (180°). This indicates that, on average, mitosis occurs in the transition zone between the contractile front and the expanding tissue interior.

Figure 5 B shows the temporal evolution of the mean divergence within the central ROI, averaged across all 90 analyzed events on a temporal scale of 100 min before and after mitosis onset. The data shows a clear sign reversal from negative divergence before mitosis start to positive values after mitosis, indicating a distinct cellular contraction before division followed by local expansion. The analysis of temporal mean values before and after mitosis start in the surrounding segmented ROIs, shown in Fig. 5 C, also reflects the observed sign reversal for the central ROI. However, the radial segments beyond the central cell reveal a spatially consistent divergence pattern both before and after mitosis. The segmented ROIs show contraction in the segments oriented toward the growth front (Segments 0°, 45°, and 315°), while the interior-facing segments indicate local expansion. By successfully integrating label-free deep learning detection with continuous optical flow kinematics, this framework reveals that dividing cells undergo a transient cycle of contraction and expansion, while remaining embedded within a persistent, macroscopic mechanical gradient of tissue convergence at the active growth front.

## Discussion

### Tissue-Level Spatial Organization of Growth

The combination of optical flow analysis and mitosis detection not only captures macroscopic tissue growth but also reveals how it is driven by spatially compartmentalized kinematics. In the presented microtissue, macroscopic tissue organization is governed by a circular defect – created by a spontaneous internal rupture – that progressively closes over time. The key finding of our label-free analysis in this context is the distinct spatial colocalization of elevated tissue dynamics and mitotic activity within the first ~ 75 − 100 µm of the defect edge. These spatial correlations between dynamic movement and cell proliferation show strong parallels to the spatial distribution of contractile myofibroblasts (10). Despite the differences in substrate and tissue geometry to these previous findings, we hypothesize that our ruptured microtissues exhibit a similar biological organization. Indeed, studies on our specific model confirm that myofibroblasts are heavily localized at the growth front, with cells expressing alpha-smooth muscle actin (*α*-SMA) primarily in the first 60 µm (13). Interestingly, this 60 µm range of the *α*-SMA signal correlates with the spatial scale of our dynamically measured transients and the width of the expanding region in the divergence map, indicating a direct link between the observed tissue kinematics and the accumulation of contractile myofibroblasts.

The spatial compartmentalization between the active front and the quiescent interior is further supported by the flow dynamics normal to the tissue edge. We observed a distinct positive mean normal deformation at the tissue contour, providing evidence for continuous tissue growth at the proliferative front. These mean dynamics rapidly fall off as the distance from the proliferative front increases, reflecting a sharp transition into a mechanically stable, quiescent tissue interior. As a secondary detail, at greater distances from the interface, the mean normal deformation transitions to negative values. The negative flow is likely related to the out-of-frame growth front on the right side of the tissue (see Fig. 3 D).

Beyond normal growth, the directional decomposition of our flow fields reveals a clear dominance of parallel fluctuations restricted to the active proliferative front. We hypothesize that these parallel dynamics are a direct consequence of both structural ECM organization and active mechanobiological feedback. Previous studies on this model system observed that both contractile myofibroblasts and fibronectin (Fn) fibers are highly aligned parallel to the tissue contour (13). Because cells exhibit contact guidance along these anisotropically aligned fibers and can also respond to local topographical curvature — a process known as curvotaxis (32) — cellular migration is robustly directed along the curved tissue-medium interface. Furthermore, we hypothesize that elevated fluctuations parallel to the tissue edge play a crucial role in macroscopic tissue development by facilitating active remodelling (33, 34). These localized dynamics likely reflect the continuous, tension-driven cellular rearrangements and local matrix deformations necessary for tissue growth, contrasting sharply with the steep drop in fluctuations within the relaxed core. Ultimately, the spatial correlation of contractile tissue, local dynamics, and proliferation at the growth front has a clear connection to macroscopic tissue shape and substrate geometry. It is well established that macroscopic tissue development is controlled by geometric constraints, linking tissue anchorage and curvature to local proliferation (1). The concentration of mitotic activity and tissue dynamics at the highly concave defect contour links our observed kinematics to the accelerated tissue deposition on highly concave surfaces reported by Ehrig et al. (11).

Our macroscopic tissue descriptors reveal continuous temporal correlations that reflect this geometric control: as the pore closes, the local concave curvature increases, leading to an accelerated growth of the tissue area. The influence of the underlying geometry becomes even more apparent when comparing the continuously curved circular defect with the early stage tissue geometry (Supplementary Fig. S3 and S5), where expansion signatures are notably less pronounced likely due to lower overall concavity. Importantly, this is consistent with the fundamental principle of geometry-driven orchestration, observed not only in our engineered clefts but also in broader bioengineering applications. For instance, understanding how local curvature dictates the proliferative front is relevant for optimizing the development of complex organoids (35), or for predicting closure dynamics in wound healing models (36–38). Since these macroscopic growth patterns ultimately emerge from individual cellular behaviors, we next discuss the localized kinematic footprint of single cell divisions.

### Single-Cell Mechanics and Tissue Feedback

In addition to analyzing continuous dynamics at the macroscopic tissue level, the pixel-by-pixel resolution of the optical flow method enables the investigation of localized kinematics at the single-cell level. By combining the spatial localization of mitosis events with our optical flow field divergence, we revealed distinct spatio-temporal correlations between cell division and local tissue dynamics. Specifically, we observed a contraction phase in the immediate cellular vicinity prior to actual division, followed by a subsequent local expansion phase (Fig. 5 B). We hypothesize that this sequence is driven by mitotic rounding–a well-documented process where cells retract their protrusions to adopt a circular geometry prior to cytokinesis–followed by the generation of protrusive forces during mitotic elongation and the volume increase of the two daughter cells (15). This hypothesis is visually supported by our brightfield data, where the key feature of detected mitosis events is their distinctly rounded shape immediately prior to division (shown in the representative events in Fig. 1 C). We hypothesize that this localized tissue contraction-expansion cycle reflects dynamic mechanical interactions between individual dividing cells and the ECM network in which they are embedded, an interplay that could be explicitly verified in future studies combining this framework with fluorescently labeled matrix.

Moving beyond the immediate cellular vicinity, our analysis of the segmented regional ROIs (Fig. 5 C) reveals consistent, long-range dynamic asymmetries surrounding these mitotic events. Specifically, the regional tissue facing the active growth front exhibits net expansion, whereas the region facing the mature tissue interior exhibits net contraction. This spatial asymmetry raises a fundamental question regarding causal directionality in tissue orchestration: do these macroscopic dynamic patterns emerge as a direct mechanical consequence of localized, high-frequency mitotic activity in the growth front, or are proliferating cells localized in this region because the pre-existing dynamic environment (e.g., tension gradients and parallel fluctuations) is conducive to cell division? Grounded in the established principles of reciprocal cell-ECM cross-talk, we propose that this relationship is mutually reinforcing, functioning as a continuous mechanical feedback loop. Traditionally, mapping how individual mitotic events mechanically stimulate their neighbors requires complex, multi-reporter live fluorescence imaging (39). By successfully extracting these localized biomechanical events purely from brightfield image streams, our approach provides a highly accessible, non-invasive method to map how microscale single-cell forces integrate to drive macroscopic tissue morphogenesis.

### Versatility and Limitations of the Label-Free Framework

The analysis of tissue growth using dense optical flow and YOLO-assisted object detection provides a highly versatile, label-free framework. Because it extracts multi-scale biomechanical information directly from brightfield images, it maintains the biological system in its native state, avoiding the phototoxicity and artifacts associated with fluorescent tracking. This makes the methodology highly applicable to a broad range of mechanobiological models where noninvasive, long-term monitoring is crucial. For example, this pipeline could be adapted to track the morphological development of organoids, quantify closure dynamics in advanced *in vitro* wound healing assays, or evaluate the maturation of 3D-bioprinted constructs. However, applying this framework across diverse biological systems requires careful consideration of its specific algorithmic limitations and input data requirements.

Unlike traditional Particle Image Velocimetry (PIV), which relies on spatial averaging that can obscure highly localized kinematics, dense optical flow preserves pixel-wise dynamics (40). However, the Farnebäck algorithm fundamentally assumes brightness constancy (20). This condition can be easily violated by illumination fluctuations, cellular occlusion, or the continuous inter-frame deposition of new extracellular matrix characteristic of dense microtissues. Additionally, resolving multi-scale dynamics requires a delicate balance of spatial and temporal resolution. Utilizing multiple image pyramid levels enables the algorithm to capture rapid, largescale tissue deformations, but this downscaling inevitably blurs the highly localized kinematic footprints around individual cells. Temporally, for capturing active cellular migrations and dynamic matrix deformations, a timescale of several minutes represents an effective compromise. Previous studies have successfully utilized 5-to 15-minute intervals for these processes (14, 31), confirming the suitability of the ~ 10-minute temporal resolution used in our approach. Naturally, any rapid transient movements occurring entirely between frames remain undetectable. Perhaps the most fundamental limitation of relying solely on brightfield images is the lack of biomolecular specificity; the extracted flow fields represent a continuous superposition of active cellular migration and extracellular matrix deformation. While this holistic view accurately captures overall tissue kinematics, resolving individual cell movement from matrix displacement is challenging using brightfield data alone. Conveniently, our modular optical flow framework is not restricted to brightfield images and can be readily applied to isolated fluorescence channels whenever this explicit decoupling is required.

For object detection, our YOLO-assisted approach significantly accelerated the localization of mitotic activity, reducing the overall manual annotation time by approximately 26.5%. While currently semi-automated–requiring partial manual labeling of the specific dataset to train the model– training generalized models on a number of diverse brightfield datasets could enable fully automated mitosis detection. Finally, because our input data relies on standard brightfield microscopy, the present analysis is constrained to the highresolution dynamics visible at a single 2D focal plane, which occasionally limits the tracking of deep-tissue events. Nevertheless, because our pipeline processes images on a modular, frame-by-frame basis, the methodology is fundamentally scalable. By acquiring multi-*z*-plane or true volumetric timelapse data, this label-free framework can be readily extended to capture the full 3D biomechanical complexity of developing microtissues.

## Conclusions

The proposed combination of dense optical flow motion estimation and deep learning-assisted object detection provides a powerful, label-free framework to extract continuous, multi-scale biomechanical information directly from standard brightfield microscopy. At the macroscopic level, our methodology successfully compartmentalizes the developing tissue into an active growth front and a quiescent mature interior. We demonstrate that dynamic tissue growth and remodeling are strictly localized to the highly curved growth front. This active zone is distinctly characterized by a concentration of mitotic events, highly elevated parallel fluctuations, and positive normal expansion, directly reflecting how underlying substrate geometry and concave curvature dictate the spatiotemporal orchestration of tissue growth.

At the microscopic scale, the method successfully captures the localized kinematic footprints of individual cell divisions, revealing a distinct sequence of tissue contraction and expansion in the immediate vicinity of mitotic events. Because these dynamically active, proliferating cells are predominantly localized at the growth front, our framework highlights a complex interplay connecting local cellular kinetics to emergent macroscopic tissue growth. Importantly, this spatial correlation allows for a bidirectional interpretation: it remains to be determined whether dividing cells localize at the growth front in response to the pre-existing mechanobiological environment, or if the distinct local mechanobiology–characterized by reciprocal mechanical feedback and push-and-pull forces–emerges as a direct consequence of these localized mitotic events.

While the exact causal hierarchy of how these cellular feedback interactions ultimately determine overall tissue development cannot be fully resolved by label-free kinematic analysis alone, the presented methodology serves as a non-destructive tool for dynamic tissue evaluation. A key challenge for future investigations will be to explicitly decouple individual cell migration from extracellular matrix deformation. Resolving this distinction will further illuminate how microscopic single-cell forces collectively emerge to drive the macroscopic maturation of engineered tissues. To facilitate such investigations and support the broader mechanobiology community, the complete analysis framework is provided as a ready-to-use, open-source tool. We highly encourage researchers to apply, modify, and expand upon this modular pipeline to extract continuous kinematic insights across diverse biological model systems.

## Supporting information

Supplementary Movie 2: Animation of deformation direction over time

Supplementary Movie 1: Animation of deformation magnitude over time

## Code and Data Availability

To support reproducibility and facilitate further research, the complete computational pipeline developed in this study–including dense optical flow estimation, spatial mitosis analysis, and geometric tissue analysis–is provided as a modular, open-source tool. The Python source code, along with instructions for implementation and modification, is publicly available on GitHub at: https://github.com/LennardFastabend/microtissue-flow-mitosis

Additionally, due to file size constraints, the raw brightfield image time-series and manually curated YOLOv5 mitosis labels required to reproduce the main findings are hosted on Zenodo. The dataset can be accessed via DOI: https://doi.org/10.5281/zenodo.21238557.

## Author Contributions

**Conceptualization:** All authors.

**Methodology:** KLF, TVT, MCB.

**Software:** KLF, RC.

**Formal Analysis:** KLF.

**Investigation:** KLF, TVT, MCB.

**Visualization:** KLF, PK.

**Writing – Original Draft:** KLF, PK.

**Writing – Review & Editing:** All authors.

**Supervision:** VV, PK.

**Project Administration:** VV, PK.

**Funding Acquisition:** VV, PK.

## Acknowledgments

Financial support from the following funding agencies is gratefully acknowledged: Swiss NSF National Center of Competence in Research (NCCR) for Molecular Systems Engineering [grants NCH 1734, NCH 1791], SNF Swiss National Science Foundation [grants 204345, 175839, 219990]. BRCCH Botnar Research Center for Child Health Multi-Investigator Project 2020.

## Declaration of Generative AI and AI-assisted technologies in the writing process

During the preparation of this work, the authors used ChatGPT (OpenAI) and Gemini (Google) for coding assistance, DeepL (DeepL SE) for translation, and Gemini to improve the readability and language of the manuscript. After using these tools, the authors critically reviewed and edited the content as needed and take full responsibility for the final content of the publication.

## Supplementary Note 1: YOLO-Assisted Labeling

**Table S1.** Representative time effort of steps in the YOLO-assisted workflow. A timeseries of 122 frames was separated into training, validation, and inference data. Training and validation frames are annotated manually and used to finetune the pretrained YOLO model. The remaining unlabeled frames are annotated by the YOLO model and corrected manually afterwards. Finally the manual labels and YOLO-assisted labels are combined and checked for consistency.

|  |  |  |  |  |  |  |
| --- | --- | --- | --- | --- | --- | --- |
| Process Step | Manual annotation of 48 training frames | Manual annotation of 12 validation frames | YOLO training | Inference on 62 unlabeled frames | Correction of YOLO annotations | Final consistency check |
| Duration | 48:30 min | 12:00 min | 7 min | 33 s | 16:20 min | 5:20 min |

**Table S2.** YOLOv5 training hyperparameters and augmentation settings.

| Optimizer & Learning Rate |  | Loss Gains & Thresholds |  |
| --- | --- | --- | --- |
| Hyperparameter | Value | Hyperparameter | Value |
| lr0 (Initial LR) | 0.01 | box (Box loss gain) | 0.025 |
| lrf (Final LR) | 0.005 | cls (Cls loss gain) | 0.3 |
| momentum | 0.937 | cls_pw (Cls BCELoss) | 1.0 |
| weight_decay | 0.0005 | obj (Obj loss gain) | 0.7 |
| warmup_epochs | 3.5 | obj_pw (Obj BCELoss) | 1.0 |
| warmup_momentum | 0.8 | iou_t (IoU threshold) | 0.20 |
| warmup_bias_lr | 0.1 | anchor_t | 4.0 |
|  |  | fl_gamma | 0.0 |
| Data Augmentation (Probability or Magnitude) |  |  |  |
| hsv_h (Hue) | 0 | hsv_s (Saturation) | 0 |
| hsv_v (Value) | 0.5 | degrees (Rotation) | 20 |
| translate | 0.1 | scale | 0.8 |
| shear | 0.0 | perspective | 0.0 |
| flipud (Up-Down) | 0.5 | fliplr (Left-Right) | 0.5 |
| mosaic | 1.0 | mixup | 0 |
| copy_paste | 0 |  |  |

**Fig. S1.**
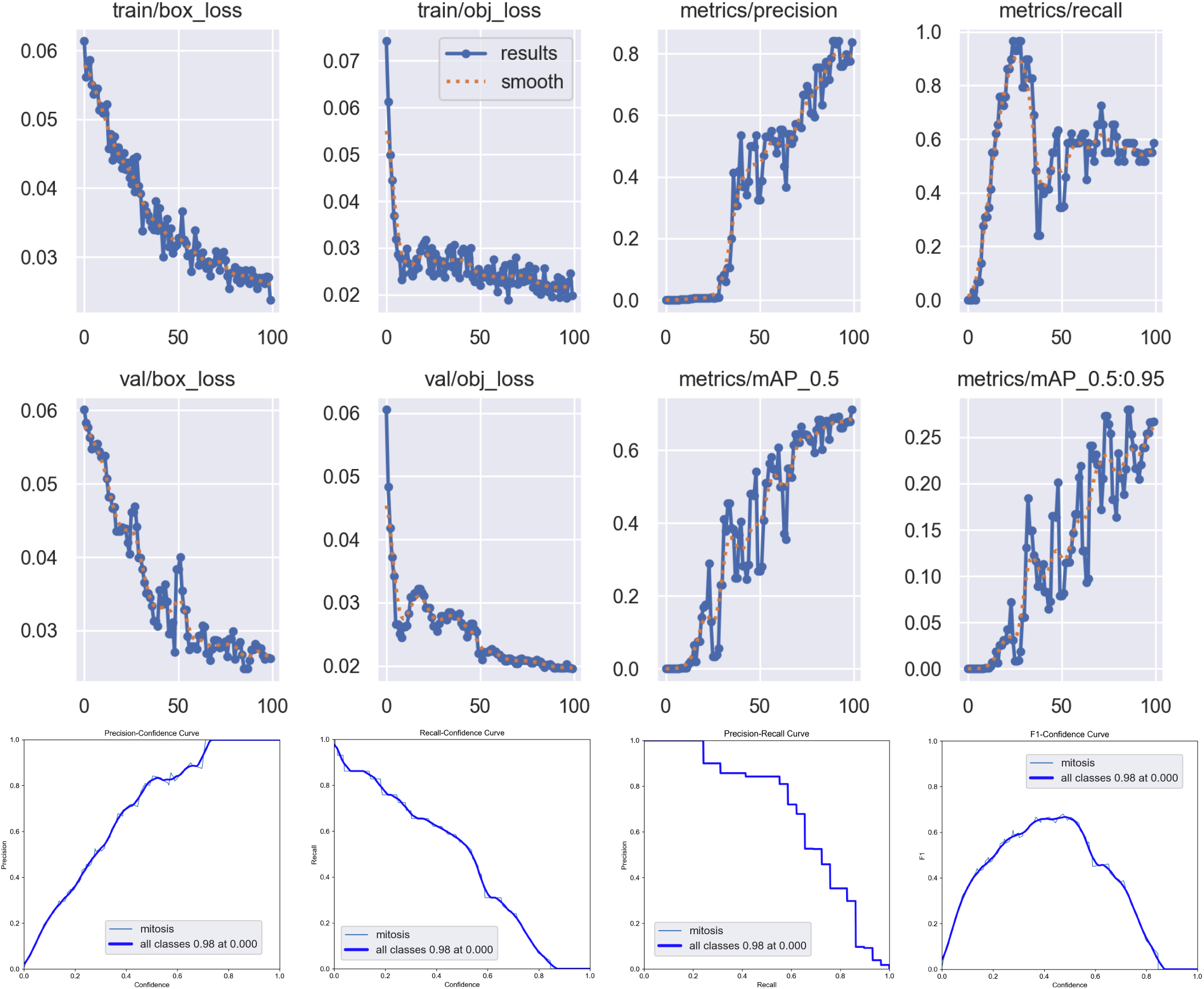
YOLO training results on manually labeled frames. 122 total frames are split into unlabeled and labeled frames at random. 60 labeled frames are used to finetune the pretrained YOLO model yolov5s over 100 epochs. Of the 60 labeled frames 48 frames serve as training data and 12 frames are used as validation data.

## Supplementary Note 2: Segmentation

**Fig. S2.**
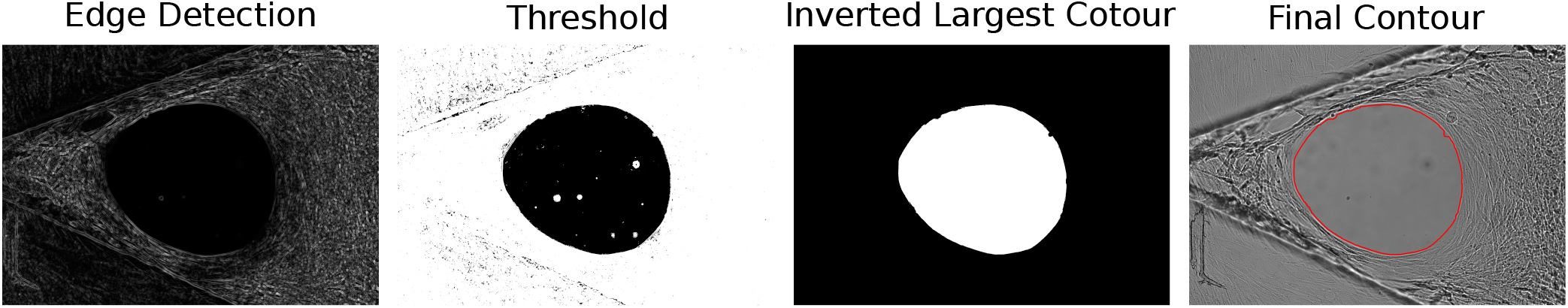
Workflow for automated tissue segmentation. A Sobel filter is applied for edge detection, followed by spatial smoothing of the gradient image. An intensity threshold is then implemented to generate a preliminary binary mask of the tissue contour. This mask is subsequently inverted, retaining only the largest connected contour. Finally, the tissue contour is smoothed and eroded to eliminate boundary offsets introduced during the initial filtering steps.

**Table S3.** Parameters used for the image segmentation pipeline.

| Parameter | Value | Description |
| --- | --- | --- |
| front_sobel_ksize | 31 | Kernel size for the Sobel operator used to compute the initial image gradients. |
| front_gauss_ksize | 3 | Kernel size for the Gaussian blur applied to the gradient image to suppress high-frequency noise. |
| front_gauss_sigma | 3 | Standard deviation ($\sigma$) for the Gaussian kernel, defining the smoothing intensity on the gradient field. |
| front_segmentation_th | 2 | Intensity threshold applied to the smoothed gradient image to generate the initial binary mask. |
| front_masksmoothing_ksize | 25 | Kernel size for the morphological opening (MORPH_OPEN) to remove small noise artifacts and smooth mask boundaries. |
| front_erosion_ksize | 3 | Kernel size for the morphological operation used to refine the final mask shape. |
| front_erosion_iters | 3 | Number of iterations for the shape refinement step to ensure a consistent mask boundary. |

## Supplementary Note 3: Mathematical Background of the Farnebäck Algorithm

The following overview of the algorithm follows the descriptions provided by Gunnar Farnebäck in his thesis on ‘Polynomial Expansion for Orientation and Motion Estimation’ (20). The Farnebäck algorithm is based on the polynomial expansion of a pixel neighbourhood in an image to approximate local image gradients with a two-dimensional quadratic polynomial. The local image signal *f* (*x*) is therefore modeled by the function:

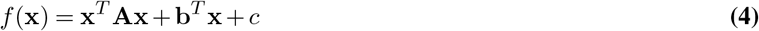

where **x** is a position in the image, **A** is a symmetric matrix, **b** a vector and *c* a scalar value. To compute the coefficients **A, b** and *c*, a normalized convolution of the local image signal with the basis functions (Eq. 5) is performed. For the calculation of the coefficients in **A** and **b** this is analogous to the calculation of the first and second-order derivatives of the local image signal. Therefore the convoluted images represent the local gradients and curvatures of the image intensity.

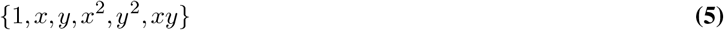

The resulting six expansion coefficients for the six base functions can be defined as:

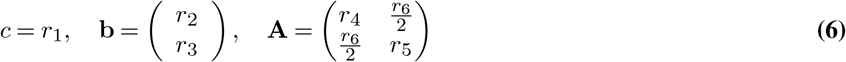

such that the signal *f* (**x**) is represented by:

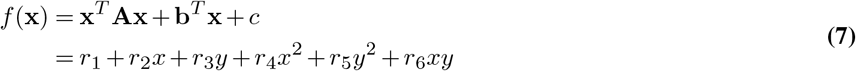

A direct comparison of the basis functions (5) with formula (7) shows that the matrix **A** contains information about the even part of the signal (without the constant offset value captured in *c*), whereas **b** captures information about the odd part of the image signal.

To understand how the Farnebäck algorithm utilizes the approximation of a neighbourhood by polynomial expansion to estimate the local displacement between two frames, it is helpful to look at the global displacement of a quadratic polynomial. Comparing the quadratic polynimial *f*_1_(**x**) with a shifted signal _2_(**x**) = *f*_1_(**x** − **d**) yields:

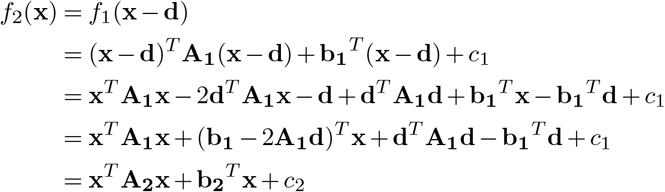

The direct comparison of the coefficients in *f*_1_ and *f*_2_ results in:

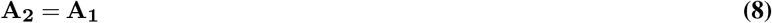

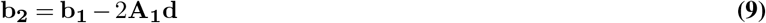

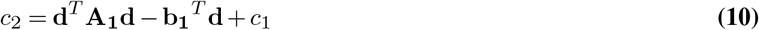

If we take equation (9) and assume that **A**_**1**_ is invertible we can rearrange it to:

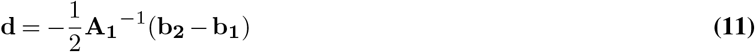

This implicates, that we can calculate the displacement of the signal from the expansion coefficients **A**_**1**_, **b**_**1**_ and **b**_**2**_.

**Table S4.**
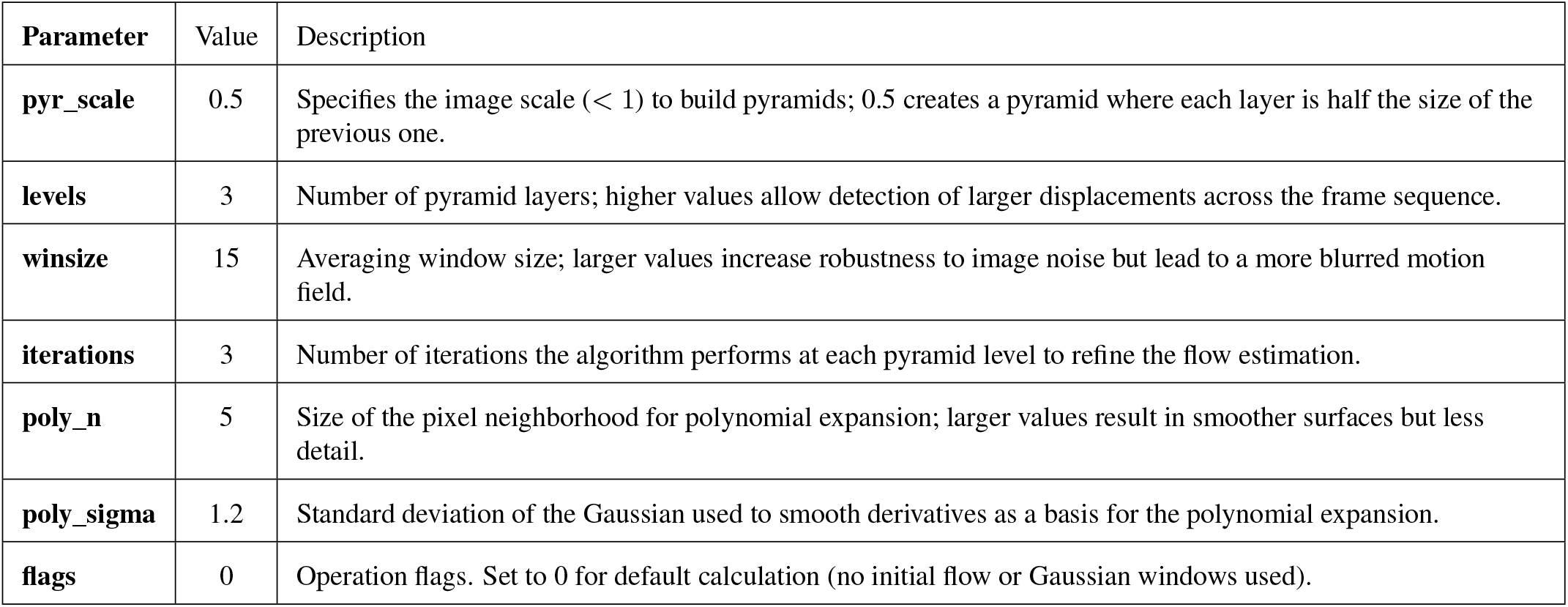
Parameters used for the Farnebäck Dense Optical Flow algorithm implementation in OpenCV with descriptions based on the documentation on the *cv2.calcOpticalFlowFarneback* function (23).

| Parameter | Value | Description |
| --- | --- | --- |
| **pyr_scale** | 0.5 | Specifies the image scale ($< 1$) to build pyramids; 0.5 creates a pyramid where each layer is half the size of the previous one. |
| **levels** | 3 | Number of pyramid layers; higher values allow detection of larger displacements across the frame sequence. |
| **winsize** | 15 | Averaging window size; larger values increase robustness to image noise but lead to a more blurred motion field. |
| **iterations** | 3 | Number of iterations the algorithm performs at each pyramid level to refine the flow estimation. |
| **poly_n** | 5 | Size of the pixel neighborhood for polynomial expansion; larger values result in smoother surfaces but less detail. |
| **poly_sigma** | 1.2 | Standard deviation of the Gaussian used to smooth derivatives as a basis for the polynomial expansion. |
| **flags** | 0 | Operation flags. Set to 0 for default calculation (no initial flow or Gaussian windows used). |

## Supplementary Note 4: Flow Field Analysis of Early Stage Tissue

**Fig. S3.**
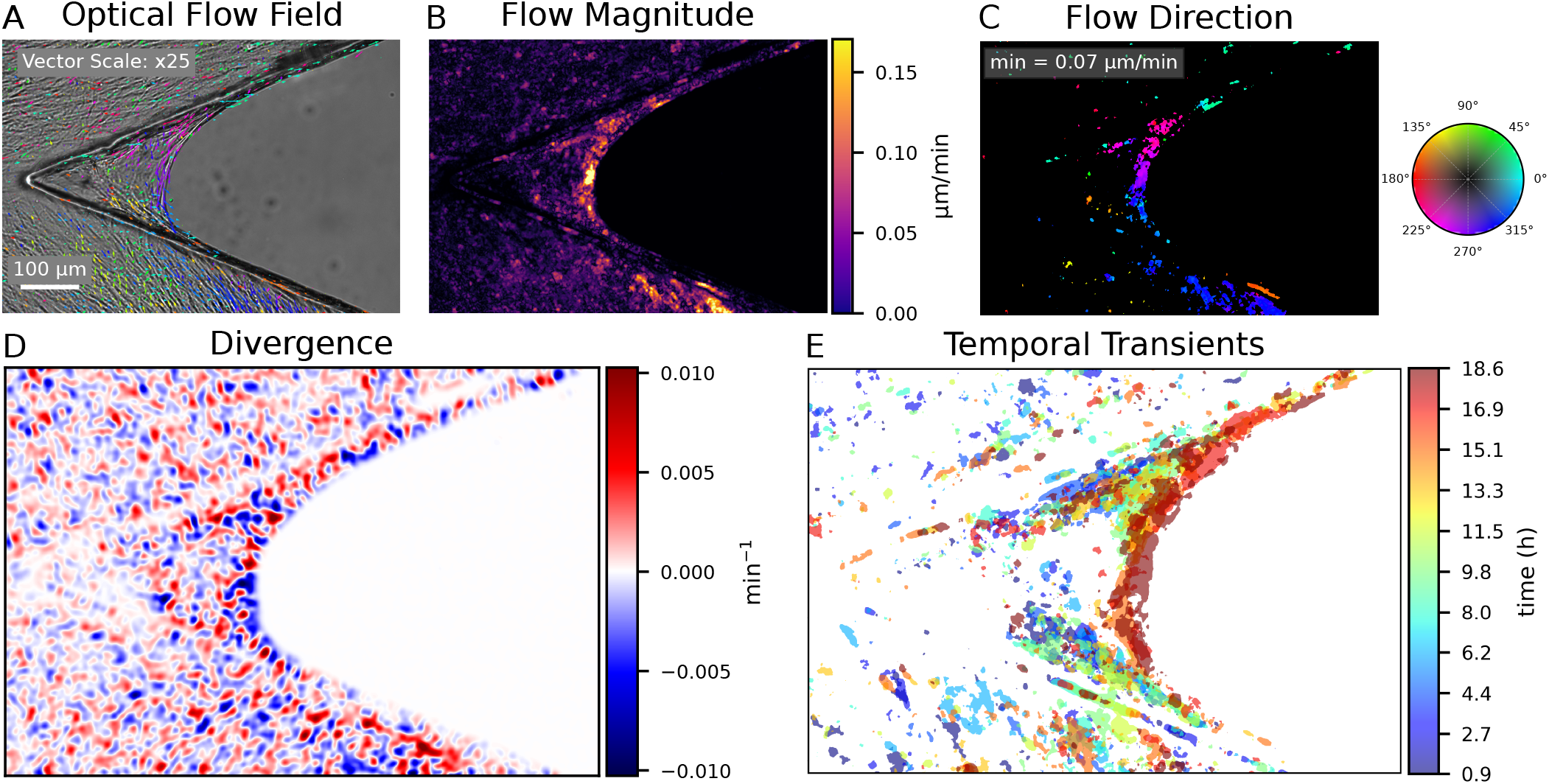
Optical flow analysis of early stage tissue. **(A)** Representative optical flow field, visualized as a vector field on the first image of the considered time window (stride: 50 px; vector scale: x25). Flow fields are averaged over 10 consecutive frames to highlight persistent long-term motion. **(B-C)** The flow magnitude (B) and direction (C) illustrate the pixel-level information contained in the flow field. The flow direction is filtered by a minimum velocity threshold of 0.07 µm/min to isolate dynamic transients. **(D)** Divergence of the optical flow field: positive divergence (sources) indicates regions of apparent tissue expansion, while negative divergence (sinks) corresponds to apparent compression. The divergence map reveals granular spatial patterns of alternating expansion and compression, highlighting flow convergence at the growth front. **(E)** Temporal evolution of dynamic transients (> 0.07 µm/min) calculated across non-overlapping time windows for the full duration of the time series, color-coded by time. The spatial distribution of the temporal transients coincides with the progressively closing pore, illustrating the spatio-temporal dynamics of the growth front.

## Supplementary Note 5: Geometric Quantification

**Fig. S4.**
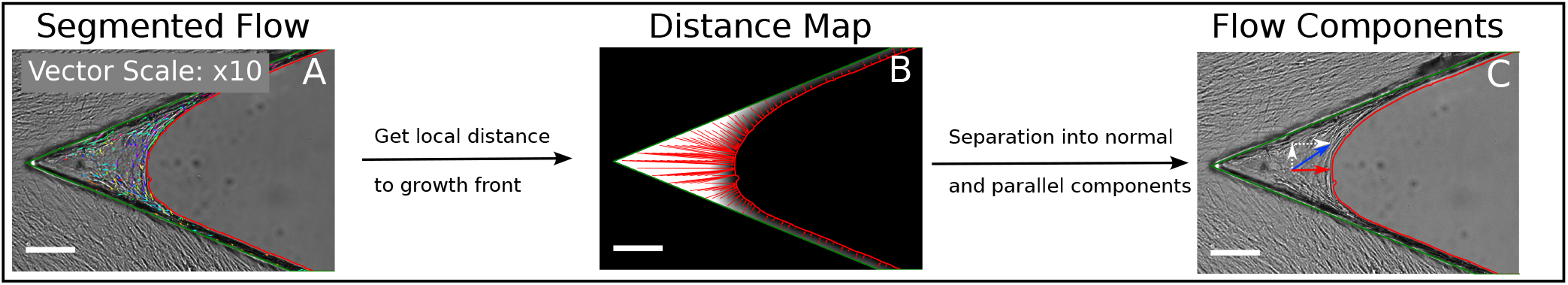
Separation of parallel and normal flow components relative to the growth front. (A) The segmented contours of the cleft edges (green) and the growth front (red) are used to mask the dynamics of tissue inside the cleft. (B) The distance map defines a vector for each pixel inside the tissue mask pointing to the closest point of the growth front contour. The underlying gray color of the distance map is defined by the length of the local distance vector. (C) The local flow vector is separated into two orthogonal components with respect to the local distance vector. The here called normal components point in the same direction as the distance vector, while the parallel component is orthogonal to the local distance vector and therefore parallel to the growth front. This component separation is performed pixel wise for each local pair of flow and distance vector and the normal and parallel deformations can be visualized as individual deformation maps

**Fig. S5.**
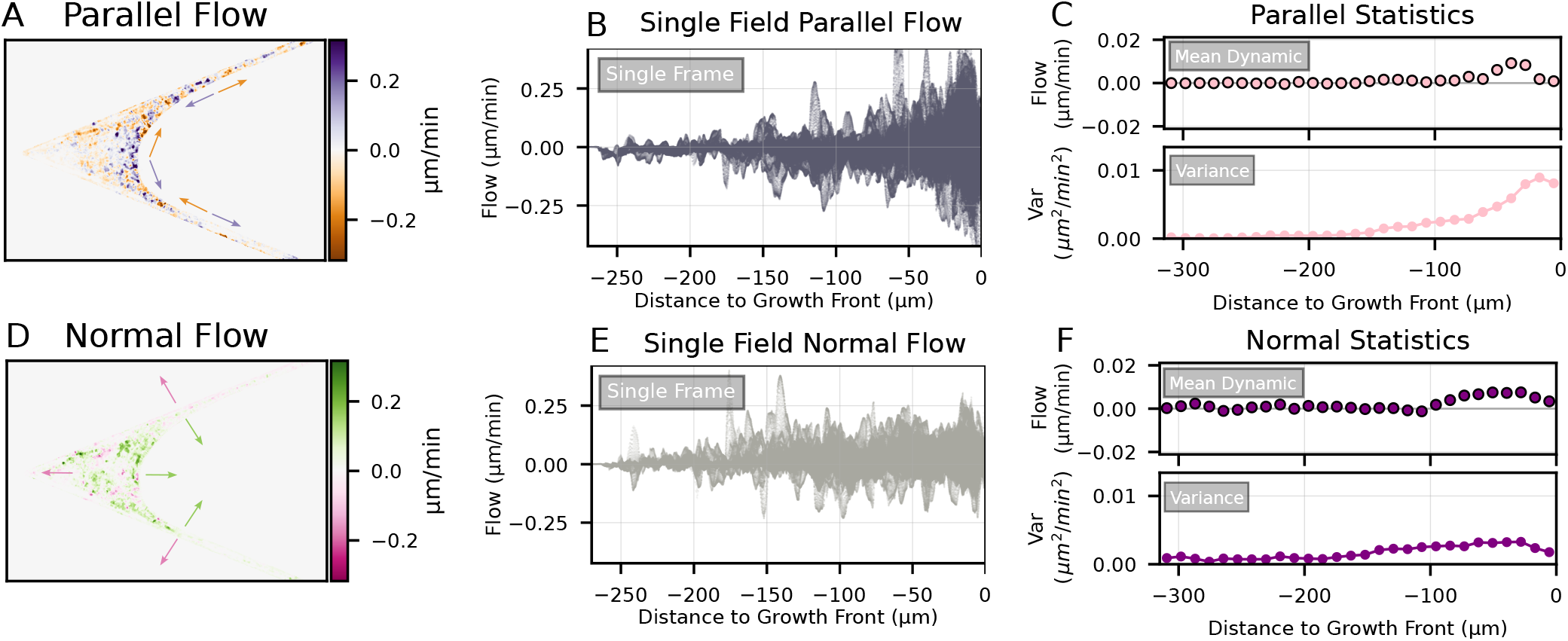
Geometric quantification of normal and parallel flow in early stage tissue. **(A,D)** Local optical flow separated into parallel (A) and normal (D) components relative to the growth front contour, visualized as scalar fields. Arrows indicate the local direction for each component at the tissue contour, corresponding to the diverging color scales. **(B,E)** Scatter plots of local magnitude against growth front distance for parallel (B) and normal (E) flow of a single field. Each point in the scatter plot corresponds to one pixel in the scalar field. **(C,F)** Temporal statistics of mean flow and variance over the entire time series, calculated in 50 px distance bins. Parallel statistics (C) reveal increased fluctuations and zero-mean flow in proximity to the growth front, while normal statistics (F) are characterized by positive mean dynamics at the growth front with consistently low variance across the entire tissue.

## Bibliography

1. Barbara Schamberger, Ricardo Ziege, Karine Anselme, Martine Ben Amar, Michał Bykowski André P. G. Castro, Amaia Cipitria, Rhoslyn A. Coles, Rumiana Dimova, Michaela Eder, Sebastian Ehrig, Luis M. Escudero, Myfanwy E. Evans, Paulo R. Fernandes, Peter Fratzl, Liesbet Geris, Notburga Gierlinger, Edouard Hannezo, Aleš Iglič, Jacob J. K. Kirkensgaard, Philip Kollmannsberger, Łucja Kowalewska, Nicholas A. Kurniawan, Ioannis Papantoniou, Laurent Pieuchot, Tiago H. V. Pires, Lars D. Renner, Andrew O. Sageman-Furnas, Gerd E. Schröder-Turk, Anupam Sengupta, Vikas R. Sharma, Antonio Tagua, Caterina Tomba, Xavier Trepat, Sarah L. Waters, Edwina F. Yeo, Andreas Roschger, Cécile M. Bidan, and John W. C. Dunlop. Curvature in biological systems: Its quantification, emergence, and implications across the scales. Advanced Materials, 35(13):2206110, 2023. doi: 10.1002/adma.202206110.

2. Celeste M. Nelson and Mina J. Bissell. Of extracellular matrix, scaffolds, and signaling: Tissue architecture regulates development, homeostasis, and cancer. Annual Review of Cell and Developmental Biology, 22(Volume 22, 2006):287–309, 2006. ISSN 1530-8995. doi: 10.1146/annurev.cellbio.22.010305.104315.

3. Jasper Foolen, Tadahiro Yamashita, and Philip Kollmannsberger. Shaping tissues by balancing active forces and geometric constraints. Journal of Physics D: Applied Physics, 49 (5):053001, ec 2015. doi: 10.1088/0022-3727/49/5/053001.

4. Jay D Humphrey, Eric R Dufresne, and Martin A Schwartz. Mechanotransduction and extracellular matrix homeostasis. Nature reviews Molecular cell biology, 15(12):802–812, 2014. doi: 10.1038/nrm3896.

5. Linda G Griffith and Melody A Swartz. Capturing complex 3d tissue physiology in vitro. Nature reviews Molecular cell biology, 7(3):211–224, 2006. doi: 10.1038/nrm1858.

6. Renjian Xie, Vaibhav Pal, Yanrong Yu, Xiaolu Lu, Mengwei Gao, Shijie Liang, Miao Huang, Weijie Peng, and Ibrahim T. Ozbolat. A comprehensive review on 3d tissue models: Biofabrication technologies and preclinical applications. Biomaterials, 304:122408, 2024. ISSN 0142-9612. doi: 10.1016/j.biomaterials.2023.122408.

7. Monika Rumpler, Alexander Woesz, John WC Dunlop, Joost T Van Dongen, and Peter Fratzl. The effect of geometry on three-dimensional tissue growth. Journal of the Royal Society Interface, 5(27):1173–1180, 2008. doi: 10.1098/rsif.2008.0064.

8. Krishna P Kommareddy, Claudia Lange, Monika Rumpler, John WC Dunlop, Inderchand Manjubala, Jing Cui, Karl Kratz, Andreas Lendlein, and Peter Fratzl. Two stages in three-dimensional in vitro growth of tissue generated by osteoblastlike cells. Biointerphases, 5(2): 45–52, 2010. doi: 10.1116/1.3431524.

9. Pascal Joly, Georg N Duda, Martin Schöne, Petra B Welzel, Uwe Freudenberg, Carsten Werner, and Ansgar Petersen. Geometry-driven cell organization determines tissue growths in scaffold pores: consequences for fibronectin organization. PloS one, 8(9):e73545, 2013. doi: 10.1371/journal.pone.0073545.

10. Philip Kollmannsberger, Cécile M. Bidan, John W. C. Dunlop, Peter Fratzl, and Viola Vogel. Tensile forces drive a reversible fibroblast-to-myofibroblast transition during tissue growth in engineered clefts. Science Advances, 4(1):eaao4881, 2018. doi: 10.1126/sciadv.aao4881.

11. S. Ehrig, B. Schamberger, C. M. Bidan, A. West, C. Jacobi, K. Lam, P. Kollmannsberger, A. Petersen, P. Tomancak, K. Kommareddy, F. D. Fischer, P. Fratzl, and John W. C. Dunlop. Surface tension determines tissue shape and growth kinetics. Science Advances, 5(9): eaav9394, 2019. doi: 10.1126/sciadv.aav9394.

12. Kai Lennard Fastabend, Cécile M. Bidan, John W. C. Dunlop, and Philip Kollmannsberger. Cortical tension links curvature to tissue growth in the cellular potts model. Phys. Rev. E, 113:024403, Feb 2026. doi: 10.1103/z152-x4l1.

13. Mario C. Benn, Simon A. Pot, Jens Moeller, Tadahiro Yamashita, Charlotte M. Fonta, Gertraud Orend, Philip Kollmannsberger, and Viola Vogel. How the mechanobiology orchestrates the iterative and reciprocal ecm-cell cross-talk that drives microtissue growth. Science Advances, 9(13):eadd9275, 2023. doi: 10.1126/sciadv.add9275.

14. Philipp Riedl and Tilo Pompe. Functional label-free assessment of fibroblast differentiation in 3d collagen-i-matrices using particle image velocimetry. Biomater. Sci., 9:5917–5927, 2021. doi: 10.1039/D1BM00638J.

15. Sungmin Nam, Yung-Hao Lin, Taeyoon Kim, and Ovijit Chaudhuri. Cellular pushing forces during mitosis drive mitotic elongation in collagen gels. Advanced Science, 8(4):2000403, 2021. doi: 10.1002/advs.202000403.

16. Cornelia Clemens, Rosa Gehring, Philipp Riedl, and Tilo Pompe. Matrix deformation and mechanotransduction as markers of breast cancer cell phenotype alteration at matrix interfaces. Biomaterials Science, 13(6):1578–1589, 03 2025. ISSN 2047-4830. doi: 10.1039/d4bm01589d.

17. Sangwoo Kim, Marie Pochitaloff, Georgina A Stooke-Vaughan, and Otger Campàs. Embryonic tissues as active foams. Nature physics, 17(7):859–866, 2021. doi: 10.1038/s41567-021-01215-1.

18. Wesley R Legant, Christopher S Chen, and Viola Vogel. Force-induced fibronectin assembly and matrix remodeling in a 3d microtissue model of tissue morphogenesis. Integrative Biology, 4(10):1164–1174, 2012. doi: 10.1039/c2ib20059g.

19. Matthew S. Hall, Rong Long, Xinzeng Feng, YuLing Huang, Chung-Yuen Hui, and Mingming Wu. Toward single cell traction microscopy within 3d collagen matrices. Experimental Cell Research, 319(16):2396–2408, 2013. ISSN 0014-4827. doi: 10.1016/j.yexcr.2013.06.009. Special Issue: Cell Motility and Mechanics.

20. Gunnar Farnebäck. Polynomial Expansion for Orientation and Motion Estimation. PhD thesis, Linköping University, Linköping, Sweden, 2002.

21. Joseph Redmon, Santosh Divvala, Ross Girshick, and Ali Farhadi. You only look once: Unified, real-time object detection. arXiv preprint arXiv:1506.02640, 2016.

22. Pietro Speziale, Livia Visai, Simonetta Rindi, and Antonella Di Poto. Purification of human plasma fibronectin using immobilized gelatin and arg affinity chromatography. Nature protocols, 3(3):525–533, 2008. doi: 10.1038/nprot.2008.12.

23. Gary Bradski. The opencv library. Dr. Dobb’s Journal of Software Tools, 25(11):120–123, 2000.

24. Glenn Jocher. Ultralytics yolov5 (version 7.0). GitHub repository, 2020. URL: https://github.com/ultralytics/yolov5, xDOI: 10.5281/zenodo.3908559.

25. Rahima Khanam and Muhammad Hussain. What is yolov5: A deep look into the internal features of the popular object detector. arXiv preprint arXiv:2407.20892, 2024.

26. Tzutalin. Labelimg. GitHub repository, 2015. URL: https://github.com/tzutalin/labelImg.

27. Cécile M. Bidan, Krishna P. Kommareddy, Monika Rumpler, Philip Kollmannsberger, Yves J. M. Bréchet, Peter Fratzl, and John W. C. Dunlop. How linear tension converts to curvature: Geometric control of bone tissue growth. PLOS ONE, 7(5):1–11, 05 2012. doi: 10.1371/journal.pone.0036336.

28. Cécile M. Bidan, Krishna P. Kommareddy, Monika Rumpler, Philip Kollmannsberger, Peter Fratzl, and John W. C. Dunlop. Geometry as a factor for tissue growth: Towards shape optimization of tissue engineering scaffolds. Advanced Healthcare Materials, 2(1):186–194, 2013. doi: 10.1002/adhm.201200159.

29. Meghan K. Driscoll, Colin McCann, Rael Kopace, Tess Homan, John T. Fourkas, Carole Parent, and Wolfgang Losert. Cell shape dynamics: From waves to migration. PLOS Computational Biology, 8(3):1–10, 03 2012. doi: 10.1371/journal.pcbi.1002392.

30. Cécile M Bidan, Philip Kollmannsberger, Vanessa Gering, Sebastian Ehrig, Pascal Joly, Ansgar Petersen, Viola Vogel, Peter Fratzl, and John WC Dunlop. Gradual conversion of cellular stress patterns into pre-stressed matrix architecture during in vitro tissue growth. Journal of The Royal Society Interface, 13(118):20160136, 2016. doi: 10.1098/rsif.2016.0136.

31. Pardis Pakshir, Moien Alizadehgiashi, Boaz Wong, Nuno Miranda Coelho, Xingyu Chen, Ze Gong, Vivek B Shenoy, Christopher A McCulloch, and Boris Hinz. Dynamic fibroblast contractions attract remote macrophages in fibrillar collagen matrix. Nature communications, 10(1):1850, 2019. doi: 10.1038/s41467-019-09709-6.

32. Laurent Pieuchot, Julie Marteau, Alain Guignandon, Thomas Dos Santos, Isabelle Brigaud, Pierre-François Chauvy, Thomas Cloatre, Arnaud Ponche, Tatiana Petithory, Pablo Rougerie, et al. Curvotaxis directs cell migration through cell-scale curvature landscapes. Nature communications, 9(1):3995, 2018. doi: 10.1038/s41467-018-06494-6.

33. John Devany, Daniel M. Sussman, Takaki Yamamoto, M. Lisa Manning, and Margaret L. Gardel. Cell cycle–dependent active stress drives epithelia remodeling. Proceedings of the National Academy of Sciences, 118(10):e1917853118, 2021. doi: 10.1073/pnas.1917853118.

34. James W. Spurlin, Michael J. Siedlik, Bryan A. Nerger, Mei-Fong Pang, Sahana Jayaraman, Rawlison Zhang, and Celeste M. Nelson. Mesenchymal proteases and tissue fluidity remodel the extracellular matrix during airway epithelial branching in the embryonic avian lung. Development, 146(16):dev175257, 08 2019. ISSN 0950-1991. doi: 10.1242/dev.175257.

35. F. Max Yavitt, Bruce E. Kirkpatrick, Michael R. Blatchley, Kelly F. Speckl, Erfan Mohagheghian, Radu Moldovan, Ning Wang, Peter J. Dempsey, and Kristi S. Anseth. In situ modulation of intestinal organoid epithelial curvature through photoinduced viscoelasticity directs crypt morphogenesis. Science Advances, 9(3):eadd5668, 2023. doi: 10.1126/sciadv.add5668.

36. Andrea Ravasio, Ibrahim Cheddadi, Tianchi Chen, Telmo Pereira, Hui Ting Ong, Cristina Bertocchi, Agusti Brugues, Antonio Jacinto, Alexandre J Kabla, Yusuke Toyama, et al. Gap geometry dictates epithelial closure efficiency. Nature communications, 6(1):7683, 2015. doi: 10.1038/ncomms8683.

37. Wang Jin, Kai-Yin Lo, Shih–En Chou, Scott W. McCue, and Matthew J. Simpson. The role of initial geometry in experimental models of wound closing. Chemical Engineering Science, 179:221–226, 2018. ISSN 0009-2509. doi: 10.1016/j.ces.2018.01.004.

38. Min Bao, Jing Xie, Aigars Piruska, Xinyu Hu, and Wilhelm T. S. Huck. Microfabricated gaps reveal the effect of geometrical control in wound healing. Advanced Healthcare Materials, 10(4):2000630, 2021. doi: 10.1002/adhm.202000630.

39. Marina Uroz, Sabrina Wistorf, Xavier Serra-Picamal, Vito Conte, Marta Sales-Pardo, Pere Roca-Cusachs, Roger Guimerà, and Xavier Trepat. Regulation of cell cycle progression by cell–cell and cell–matrix forces. Nature cell biology, 20(6):646–654, 2018. doi: 10.1038/s41556-018-0107-2.

40. Tianshu Liu, Ali Merat, MHM Makhmalbaf, Claudia Fajardo, and Parviz Merati. Comparison between optical flow and cross-correlation methods for extraction of velocity fields from particle images. Experiments in Fluids, 56(8):166, 2015.

